# The directionality of collective cell delamination is governed by tissue architecture and cell adhesion in a *Drosophila* carcinoma model

**DOI:** 10.1101/2024.10.08.617152

**Authors:** Marta Mira-Osuna, Steffen Plunder, Eric Theveneau, Roland Le Borgne

## Abstract

Carcinomas originate in epithelia and are the most frequent type of cancers. Tumor progression starts with the collective delamination of live tumors out of the epithelium, and requires modulation of cell adhesion. Yet, it remains elusive exactly how remodeling of epithelial cell-cell and cell-extracellular matrix contacts contribute to metastasis onset and delamination directionality. We used *Drosophila melanogaster* larval eye disc to study cooperative oncogenesis and determine the contribution of bi- and tricellular septate junction (SJ) components to tumor progression in flat and pseudostratified epithelia. We reveal that loss-of-function of septate junction components alone can promote cell death whereas synergic interaction with oncogenic Ras triggers the collective delamination of living tumorigenic cells. Spatiotemporal analyses reveal apical and basal delamination processes differ in terms of cell identity, cell polarity and remodeling of cell-cell and cell-ECM contacts. Using a combination of *in vivo* and *in silico* approaches, we report that SJ-depleted *Ras^V12^* tumors in pseudostratified epithelia trigger tissue folding or basal collective delamination regardless of the order in which cell contacts are remodeled. In striking contrast, SJ-depleted *Ras^V12^* tumors formed in flat, squamous epithelia, first undergo apical constriction and acquire a dome-like shape. Tumors are enriched in adhesion molecules and form an apical neck at the interface of contact between mutant cells and wild-type neighbors. Concomitant to cytoskeleton remodeling, tumors emit apical protrusions in the lumen and progressively delaminate through the apical neck while remaining cohesive. Our study reveals that tissue architecture and changes in cell adhesion drive the directionality of collective delamination of neoplastic tumors out of an epithelium.

## Introduction

Epithelial tissues compartmentalize the organism into organs, multicellular rearrangements carrying out highly specialized functions. Epithelia are composed of one or multiple layers of polarized epithelial cells in which apico-basolateral polarity modules are spatially restricted, creating plasma membrane domains to which junctional complexes are localized (Buckley & St Johnston, 2022; Campanale et al., 2017; Flores-Benitez & Knust, 2016; Pickett et al., 2019; Tepass, 2012; Vasquez et al., 2021). At the interface of contact between two or three adjacent cells, bicellular and tricellular junctions form, respectively. In invertebrates like *Drosophila*, adherens junctions are formed by E-cadherin (E-cad)-Catenins complexes and are localized at the apical domain. Adherens junctions mediate intercellular adhesion and are linked to the actomyosin cytoskeleton, mechanically coupling neighboring cells together and establishing the mechanical barrier of the tissue. Septate junctions (SJ), considered somewhat analogous to the vertebrate tight junctions, are located at the basolateral domain and comprise over 30 biochemically diverse proteins, of which core-complex components include Coracle (Cora) and Nervana 2 (Nrv2). SJ regulate selective permeability across the epithelium upon establishment of a mature paracellular diffusion barrier, which becomes functional in late-embryogenesis (Baum & Georgiou, 2011; Guillot & Lecuit, 2013; Harris & Tepass, 2010; Izumi & Furuse, 2014; Oshima & Fehon, 2011). Loss-of-function of bicellular SJ core-components *nrv2* and *cora* in early embryonic stages results in embryonic lethality, suggesting bicellular SJ may contribute to epithelia homeostasis beyond their barrier functions (Genova & Fehon, 2003; Hall & Ward, 2016; Paul et al., 2003). In recent years, tricellular junctions have emerged as essential regulators of epithelia homeostasis across metazoans. Tricellular junctions function as hubs for mechanochemical signaling throughout development, acting as spatial landmarks for spindle orientation during cell division, sites for epithelia integration in cell migration and regulating cell intercalation during tissue convergent-extension (Finegan et al., 2019; Higashi et al., 2016; Ikawa et al., 2023; Ventura et al., 2022; Wang et al., 2018). Tricellular SJ form at the level of SJ and by three vertex-restricted proteins: the proteolipid protein M6, the triple-repeat protein Anakonda (Aka), and the transmembrane protein Gliotactin (Gli). Loss-of-function of *M6* or *aka*, upstream regulators of tricellular SJ assembly, results in early embryo lethality (Byri et al., 2015; Zappia et al., 2011), at stages 13, when the permeability barrier is not yet mature and the embryo undergoes intensive morphogenesis.

At the interface of contact between cells and the extracellular matrix (ECM), focal adhesions mediate mechanotransduction and signaling between cells and the extracellular matrix via heterodimeric integrin receptors coupled to the basal actin cytoskeleton. The ability of cells to coordinate their behavior in response to intrinsic genetic programs and extrinsic stimuli drives tissue-scale organizations and preserves homeostasis. Across development, the orchestration of novel multicellular rearrangements during cell division, intercalation and extrusion, requires a tightly controlled cross-talk between polarity cues, changes in adhesiveness and actomyosin-based contractility and force generation (Mira-Osuna & Le Borgne, 2024). In turn, mechanochemical cues converge at the cell interface where cell junctions must be robust, to sustain mechanical tension and prevent tissue rupture. At the same time and paradoxically, cell junctions must remain plastic, to modulate cell adhesion and packing. Junctional remodeling comprises the reinforcement, *de novo* assembly and disassembly of junctional complexes to enable tissue integrity maintenance in developing and mature epithelia.

Proteins of the rat sarcoma viral oncogene family (Ras) are involved in key cellular processes during development, from proliferation to differentiation, and appear mutated in ∼19% of patients with cancer worldwide, in human cancers with some of the highest mortality rate (Prior et al., 2020). The human oncogenes *K-Ras*, *H-Ras* and *N-Ras* have only one homologue in *Drosophila,* the oncogene Ras1 (Ras^85D^) (Simon et al., 1991). Ectopic expression of overactivated *Ras1* (*Ras^G12V^* also called *Ras^V12^*) causes developmental defects and promotes tumorigenic growth (Duchek & Rorth, 2001; Karim & Rubin, 1998; Yang & Baker, 2003). Metastasis starts with the deviation of tumors from the cell layer in a process known as epithelial-to-mesenchymal transition (EMT) (Yang et al., 2020). This first step in tumor progression is critical and the directionality in which cells leave an epithelium has a major impact on cell fate, tumor invasiveness, and ultimately, survival. Tissue-clearance processes, namely cell competition, prevent tumor initiation by inducing caspase-activation and extrusion of potentially hazardous cells (Cong & Cagan, 2024). Cell extrusion occurs apically in vertebrates and basally in invertebrates, and extruded cell ultimately die (Marinari et al., 2012; Monier et al., 2015; Rosenblatt et al., 2001; Teng et al., 2017; Villars et al., 2022). However, oncogenic mutations like *Ras^V12^* block cell death and can synergize with other genetic insults to revert the directionality of epithelial deviation and promote tumor progression (Dillard et al., 2021; Wu et al., 2010). As a result, tumors undergo delamination, deviating collectively and alive out of the epithelium, heightening tumor malingency (Gudipaty & Rosenblatt, 2017; Ohsawa et al., 2018). The molecular regulators governing the directionality of this three-dimensional rearrangement start to be elucidated (Slattum et al., 2009; Tamori et al., 2016; Villars et al., 2022). However, how this machinery is orchestrated together with junctional rearrangements, such as the dismantling of junctional complexes, remains poorly understood. Constitutive activation of MyoII in *Ras^V12^* tumors promotes basal deviation from the epithelium (Dunn etal., 2018). Null mutation of the tricellular septate junction component M6 synergizes with *Ras^V12^* and activates Rho-1 signaling. Subsequent remodeling of the cytoskeleton network is coupled to adherens junctions remodeling via Canoe (Manning et al., 2019; Sawyer et al., 2009; Yu & Zallen, 2020) and promotes tumor deviation both apically and basally (Dunn et al., 2018). Whether this is exclusive to M6, or can be a broader function of SJ in preserving the epithelial barrier remains unclear. The body of evidence highlighting bicellular SJ involvement in embryonic morphogenesis, M6 in carcinogenesis and the recently described interplay between bicellular and tricellular SJ, suggest SJ may contribute to epithelia homeostasis beyond paracellular diffusion barrier functions. However, how SJ integrity contributes to epithelial homeostasis in the context of tumor progression remains unknown.

To explore the potential role SJ play in tumor progression, we make use of the larval eye-antenna imaginal disc of *Drosophila melanogaster,* a model in cancer biology used to characterize oncogenic Ras interaction with additional mutations in which hallmarks of human cancers are recapitulated (Dillard et al., 2021; Miles et al., 2011; Pagliarini & Xu, 2003; Vidal & Cagan, 2006). The immature visual organ of *Drosophila* is composed of two cell layers: an outer flat epithelium (peripodial epithelium, PE) and an inner pseudostratified epithelium (disc proper, DP), separated by a lumen and facing each other through their apical domains. During larval development, the pseudostratified epithelium undergoes neurogenesis and terminal differentiation is mediated by cell-cell interactions, independently of cell linage. The forefront of the posterior-to-anterior wave of differentiation is called the morphogenetic furrow (MF) and can be observed as a transient indentation in the pseudostratified epithelium (Wolff & Ready, 1991). From larval stage L3 stage (L3 stage) until 24h after puparium formation, the initial pool of equipotent progenitors is sequentially specified in neuronal photoreceptors or accessory cells and patterned into an ommatidia lattice (Cagan & Ready, 1989; Wolff & Ready, 1991). Eye development culminates with the formation of a neuro-crystalline ommatidia lattice, a functional visual organ. In this study, we reveal how loss of SJ integrity in synergy with oncogenic Ras triggers apical and basal collective delamination of living cells in the *Drosophila* larval eye, the onset of metastasis and first step in tumor progression. Using spatiotemporal analyses of fixed specimen and mathematical modeling, we find that directionality of collective delamination is governed by tissue architecture and changes in cell adhesion. By developing a minimal theoretical model of collective delamination, we show that flat epithelia are highly sensitive to order in which junctional complexes are remodeled, while pseudostratified epithelia are not, and in both cases mutant cells require preferential adhesion to deviate collectively. We find, both *in silico* and *in vivo* that mutant cells in the flat epithelium exclusively undergo apical collective delamination. Conversely, mutant cells in the pseudostratified epithelium undergo collective cell movements, forming tissue folding and rosettes, the latter resulting from basal collective delamination. Our results unveil a novel role for SJ in preventing tumor progression and provide an understanding on how collective delamination is governed by tissue intrinsic features and cell adhesion to deviate living cells out of an epithelium, the first event in tumor progression.

## Results

### Loss of septate junction integrity in larval eye-antennal disc epithelium induces caspase activation and cell extrusion

First, we studied whether loss of SJ integrity in postembryonic epithelia is detrimental for cell survival. We used the larval eye-antennal imaginal disc and generated genetically heterozygote tissues containing wild-type cells and clones of mutant cells homozygote for recessive lethal SJ alleles. The eye disc is a developing organ that undergoes neurogenesis from L3 until 24h after pupariation (Ready et al., 1976).The morphogenetic furrow (MF) is the forefront of differentiation and can be identified as the transient apical constriction and indentation of the pseudostratified epithelium (Fig. 1A, B). The MF progresses across the eye field in a posterior-to-anterior fashion. Cells anterior to the MF are equipotent and become progressively patterned into and ommatidia lattice, posterior to the MF (Wolff & Ready, 1991). In cells from the flat and in the pseudostratified epithelium, adherens junctions main component E-cadherin (E-cad) is located apical to the bicellular SJ core-complex component Cora (Fig. 1A) and tricellular SJ components Aka and Gliotactin (Gli) (Fig. S1A), as described in the pupal notum (Esmangart de Bournonville & Le Borgne, 2020). Since loss of core SJ components is embryonic lethal, we used the Mosaic Analysis with Repressible Clone Marker (MARCM) system to generate genetic mosaicism in the developing organ. The tissue-restricted ey-Flipase (ey-Flp) was expressed under control of the *eyeless* promotor. In combination with the UAS/Gal4/Gal80 system, mitotic FRT-site specific recombination was induced and clones tagged with the GFP clone marker were generated (Germani et al., 2018; Luo & Wu, 2007). Previous studies reported synergic interaction with *Ras^V12^* resulted in cells deviating from the epithelium (Dunn et al., 2018). Hence, we first seeked to confirm whether loss-of-function of the M6 null allele (*M6^W186Stop^*) resulted in cells undergoing caspase-activated cell extrusion or live cell delamination.

**Figure 1.**
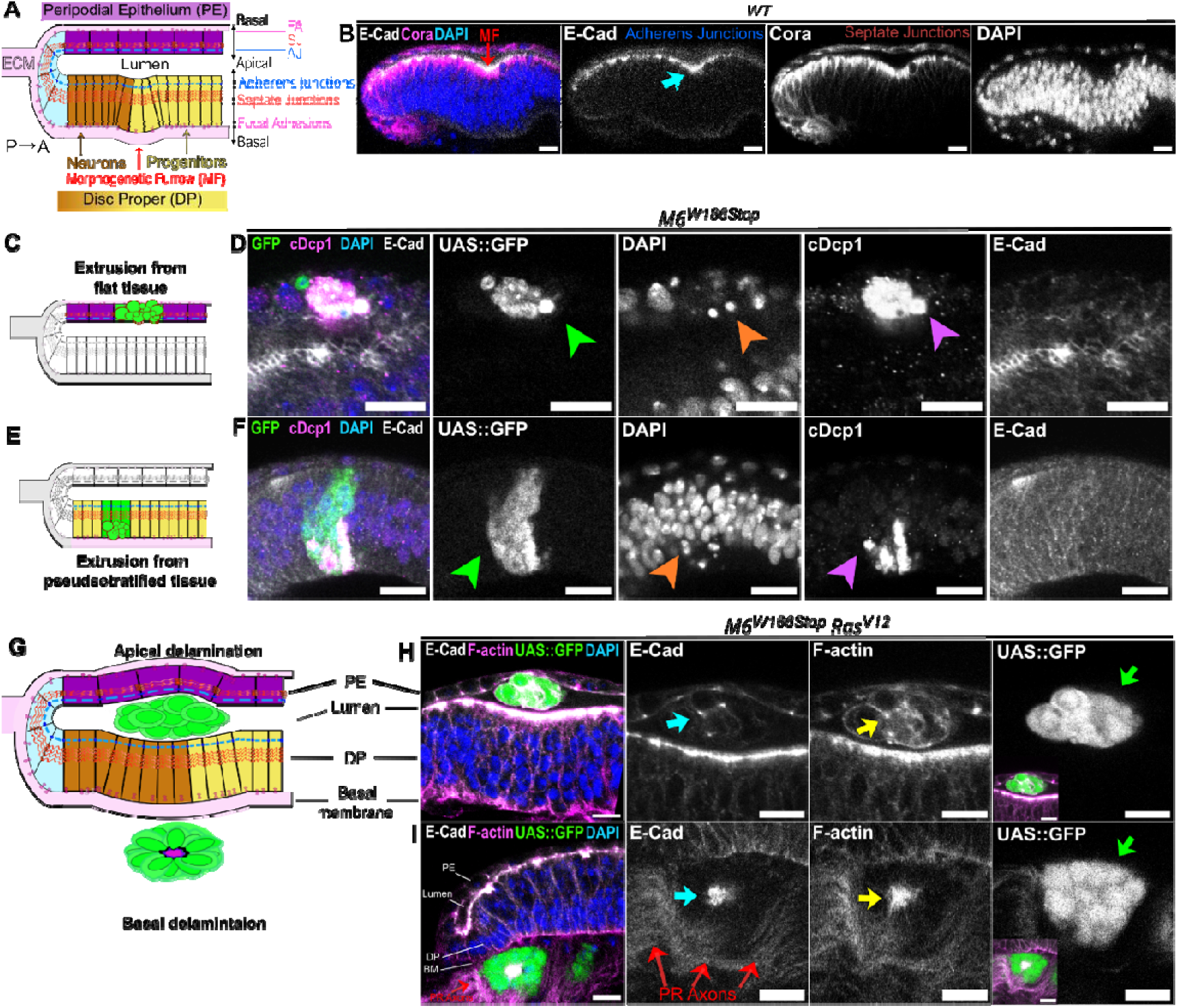
Loss of septate junction integrity in tumors triggers collective delamination of living cells from the Drosophila eye disc. (A, C, E, G) Schematic of an orthogonal section along the dorso-ventral axis of the *Drosophila* eye disc (A), apical and basal extrusion (C, E) and apical and basal delamination (G). In this and the following panels, the anterior-posterior axis is left-to-right, the peripodial epithelium (PE) in violet, undifferentiated cells in the Disc Proper (DP) in yellow, differentiated cells in the PD in brown, adherens junctions (AJ) in blue, septate junctions (SJ) in red, focal adhesions (FA) in magenta, the extracellular matrix (ECM) in light pink. Green cells show clones generated with the eyMARCM system. (B) Wild-type eye disc immunostained for E-cad, main component of adherens junctions, Coracle, core-complex SJ component, and DAPI. During development, the eye disc undergoes neurogenesis and the forefront of the anterio-posterior wave of differentiation is termed the morphogenetic furrow (MF, red arrow). MF is observed by the transient apical constriction of cells in the pseudostratified epithelium (blue arrow) and can be used as a topological landmark. (D, F) Homozygous mutant clones for M6 (*M6^W186Stop^*, green) undergo extrusion in the flat (D) and in the pseudostratified (F) epithelium. Mutant clones (green arrowheads) contain pyknotic nuclei (orange arrowheads) and are positive for the Drosophila caspase-1 (DCP-1) antibody (violet arrowheads), an upstream activator of the caspase cascade. (H, I) Homozygous mutant clones for M6 and overactivated Ras (*M6^W186Stop^Ras^V12^*, green) collectively delaminate apically in the lumen (H) and basal to the eye disc (I). Apical and basal delaminated cells are organized in clusters (green arrows), yet depending on the directionality of delamination, junctional (E-cad, blue arrow) and activated actomyosin cytoskeleton regulators markers (F-actin, yellow arrow) have striking different localization. (H) Mutant cells having undergone apical collective delamination appear in the lumen with E- cad and F-actin localized along the cell membranes. (I) Mutant cells having undergone basal collective delamination formed radial structures termed rosettes in which E-cad and F-actin localization is restricted to the central domain. Photoreceptor axons (PR axons, red arrows) can be observed in the most posterior part of the disc. The scale bars represent 10 μm. All images are single confocal optical sections with posterior to the left, anterior to the right. See also Figure S1. Figure S1. Microscopy images, related to Figures 1

Cell extrusion is the process by which junctional complexes are sequentially remodeled to seamlessly remove caspase-activated cells from the epithelium (Levayer, 2024). We observed homozygote M6 mutant cells, from here forth referred to as “clones mutant for M6” expressed the GFP-clone marker (Fig. 1D, F, green arrowheads). Clones mutant for M6 were generated in the flat (Fig. 1D) and pseudostratified epithelium, anterior (Fig. 1F) and posterior (Fig. S1B) to the MF. Clones mutant for M6 contained pyknotic nuclei (Fig. 1D, F, S1B, orange arrowheads), highly condensed spheres of DAPI-marked DNA, a sign of apoptotic activation. Caspase-activated cell extrusion is a cell competition mechanism used to eliminate polarity deficient cells from a genetically mosaic tissue when not in synergy with oncogenic Ras (Amoyel & Bach, 2014; Brumby & Richardson, 2003; Dillard et al., 2021; Igaki et al., 2009; Menéndez et al., 2010). We next studied whether clones mutant for M6 with pyknotic nuclei underwent apoptosis and cell extrusion. In the eye disc, developmentally programmed cell-death occurs during pupation in the pseudostratified epithelium, posterior to the MF, to remove the excess of unpatterned cells (Wolff & Ready, 1991). We therefore quantified cell death in the eye disc at larval stages, before developmentally programmed cell-death starts. To this aim, we made use of the marker of early caspase activation cleaved-*Drosophila* caspase-1 (cDcp1) (Fig. 1D, F, violet arrowheads). We observed over half of M6 mutant cells in the flat epithelium (Fig. 1D) and anterior to the MF in the pseudostratified epithelium (Fig. 1F) were positive for cDcp1 during L2 and early L3 larval stages. Therefore, M6 loss in flat and pseudostratified epithelia promotes caspase-activation and cell extrusion of M6-deficient cells. We next studied whether caspase-activated extrusion in M6 clones was a specific role of M6 independent of SJ, or whether it results from the disruption of SJ integrity. We studied loss-of-function mutations of bi- and tri-cellular SJ components. Single loss-of-function mutation of tricellular SJ master regulator Aka (*aka^L200^*, null allele (Byri et al., 2015)) or bicellular SJ core-component Nrv2 (*nrv2^K13315^*, hypomorphic allele (Genova & Fehon, 2003)) recapitulated the above described observations for M6. Clones of *aka* or *nrv2 mutant cells* were located in the flat (Fig. S1C, E) and pseudostratified epithelium (Fig. S1D, F) of the eye disc and were identified by the nucleus-restricted nls GFP clone marker. In these clones, nuclei were fragmented into spherical bodies, very reminiscent to pyknotic nuclei (Fig. S1C- F, orange arrowheads). Thus, mutant clones for *aka* and *nrv2* strongly resembled caspase-activated M6 clones undergoing cell extrusion. These results show loss of bi- or tri-cellular SJ components, hence SJ integrity, does not trigger detectable tissue deformation in the larval eye disc. However, SJ integrity prevents cell elimination and SJ deficient cells undergo caspase-activation and cell extrusion, possibly due to cell competition mechanisms triggered in wild-type neighbors.

### Loss of septate junction integrity synergizes with Ras overactivation to induce collective delamination

M6 loss has been observed to synergize with *Ras^V12^* and induce tumor invasion in *Drosophila* (Dunn et al., 2018). Hence, we next studied the role of bicellular and tricellular SJ components in the context of tumorigenesis to determine whether SJ integrity functions to prevent tumor progression. We induced mutant clones of a tricellular (*M6^W186Stop^*, *aka^L200^*, *gli^DV3^*) or bicellular (*nrv2^K13315^*) SJ component, in synergy with *Ras^V12^*, to make cells refractory to apoptosis. Clones with oncogenic Ras and M6 null allele underwent apical and basal collective delamination, appearing deviated in the lumen or basal to the eye disc, respectively (Fig1G-I). Moreover, we observed no sign of caspase activation in neither clones deviated apically (Fig. S2A, B) nor basally (Fig. S2C). This suggests M6-deficient tumors delaminate collectively and alive out of the epithelium. We also generated clones of cells homozygote mutant for other tricellular *(aka^L200^*, *gli^DV3^*) or bicellular (*nrv2^K13315^*) SJ components in synergy with *Ras^V12^*. Tricellular SJ-deficient tumors underwent apical (Fig. S2E) and basal collective delamination (Fig. S2F). Similarly, loss of bicellular SJ in tumors also promoted the apical (Fig. S2G) and basal (Fig. S2H) collective delamination of living cells out of the epithelium. Together, these observations show loss of SJ integrity in tumors triggers the collective delamination of cells alive out of the epithelium, both apically and basally. Multicellular organization and localization of E-cad differed between apically and basally delaminated clones. SJ-deficient *RasV^12^* cells that underwent apical collective delamination clustered in the lumen and E-cad localized along cell membranes (Fig. 1H, S2B, E, G). Conversely, SJ-deficient *RasV^12^*cells that underwent basal collective delamination formed basal rosettes, radial structures in which E-cad was restricted to the central domain (Fig1E-F). Hence, we next characterized the remodeling of junctional complexes and the actomyosin network to decipher how together they orchestrate apical and basal collective delamination.

### Tissue folding in the pseudostratified epithelium exhibits different junctional remodeling in clones anterior and posterior to the MF

We next set to decipher the sequence of events leading up to apical and basal collective delamination. We staged larval eye-antenna imaginal discs at different time-windows during larval development and performed spatiotemporal analyses of fixed specimen (see Methods). We stained for cell-cell and cell-extracellular matrix adhesion proteins, E-cad and β-integrin, respectively. We also characterized cytoskeleton remodeling using the phosphorylated regulatory light chain of non-muscle type II myosin (P-Myo) as a proxy for active MyoII and filamentous actin (F-actin). First, we studied SJ-deficient tumors located in the undifferentiated region of the pseudostratified epithelium, anterior to the MF. From late L2 until L3 stages, a gradation in the phenotype of clones was associated with clone size. Small clones were wild-type-like and did not showcase any difference in terms of junctional markers or cytoskeleton remodeling in comparison to adjacent wild-type neighbors. (Fig. 2A- C). Intermediate size clones exhibited apical constriction and basolateral spreading of E-cad and P-Myo, and remained basally anchored to the extracellular matrix via β-integrin (Fig. 2D- F). Large clones formed tissue folds and showcased apical constriction as well as basolateral spreading of E-cad and P-Myo. Strikingly, β-integrin in large clones was no longer restricted to the basal pole and instead appeared globally increased at the lateral and apical cortex (Fig. 2G-I). We next focused on SJ-deficient tumors located posterior to the MF, in the differentiated region of the pseudostratified epithelium. These clones also had a gradation in phenotype but showcased different junctional remodeling to anteriorly located clones. During L3 stages, small clones posterior to the MF underwent apical constriction concomitant with loss of E-cad basally and detachment from the extracellular matrix, evidenced by the basal loss of β-integrin and actin stress fibers (Fig. 3A-C). Bigger clones formed deeply invaginated epithelial folds and also showcased apical constriction concomitant to the basal loss of E- cad, β-integrin and F-actin. Strikingly, tissue folds were additionally enriched in β-integrin at the apical cortex (Fig. 3F). Altogether, these results indicate that synergic interaction between the loss of SJ integrity and *Ras^V12^* in the pseudostratified epithelium severely disrupts tissue architecture, triggering collective cell movements with a basal directionality. Junctional and cytoskeleton remodeling during epithelia folding is strikingly different in SJ- deficient tumors located anterior and posterior to the forefront of neural differentiation. Furthermore, differences in adhesiveness between mutant cells and their wild-type counterparts may be essential for tissue fold formation in this tissue.

**Figure 2.**
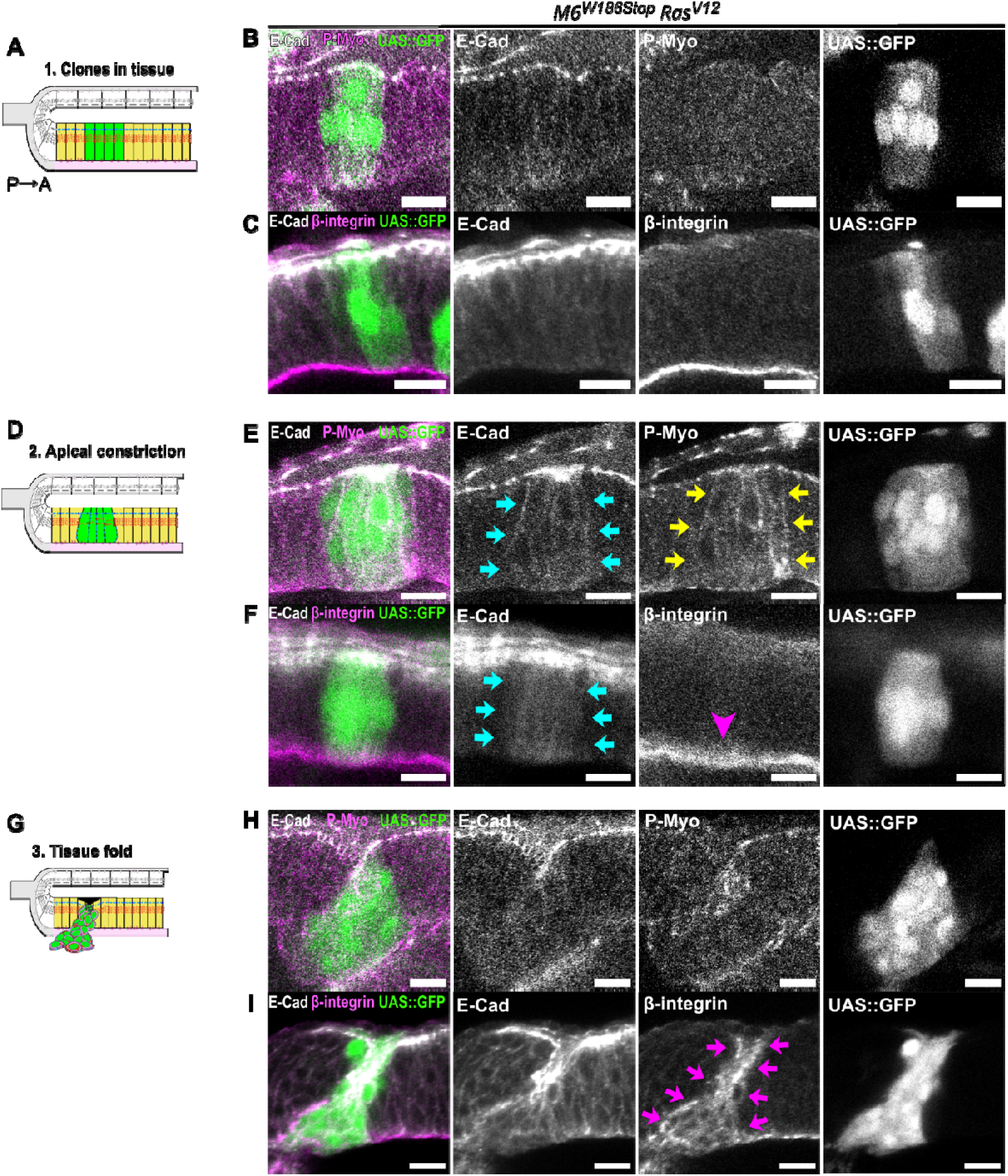
Mutant cells expressing oncogenic Ras form tissue fold anterior to the MF after apical constriction and basolateral delocalization of E-cadherin. (A, D, G) Schematic of an orthogonal section along the dorso-ventral axis of the *Drosophila* eye disc in the undifferentiated region of the pseudostratified epithelium (yellow). Mutant cells (green) initially showcase no difference to their wild-type counterparts (A), then become apically constricted (D) and finally form tissue folds (G). (B, C, E, F, H, I) Heterozygous eye disc immunostained with anti-E-cad (grey) and anti-P- Myo (B, E, H, magenta) or anti-β-integrin (C, F, I, magenta). Homozygous mutant cells for M6 (*M6^W186Stop^*) and oncogenic Ras cells express the clone marker UAS::GFP (green). (B, C) Mutant cells anterior to the MF showcase no differences in terms adherens junctions, focal adhesions or cytoskeleton remodeling. (E, F) Apical constriction of mutant cells is concomitant with E-cad basolateral spreading (blue arrows) and basolateral cytoskeleton remodeling (green arrows), while β-integrin-mediated anchoring to the extracellular matrix is preserved (pink arrowhead). (H, I) Lastly, mutant cells continue to apically constrict their surface and invaginate and exhibit basolateral spreading of E-cad and activated cytoskeleton regulators, together with basolateral spreading of β-integrin (pink arrows). The scale bars represent 10 μm. All images are single confocal optical sections with posterior to the left, anterior to the right. See also Figure S2.

**Figure 3.**
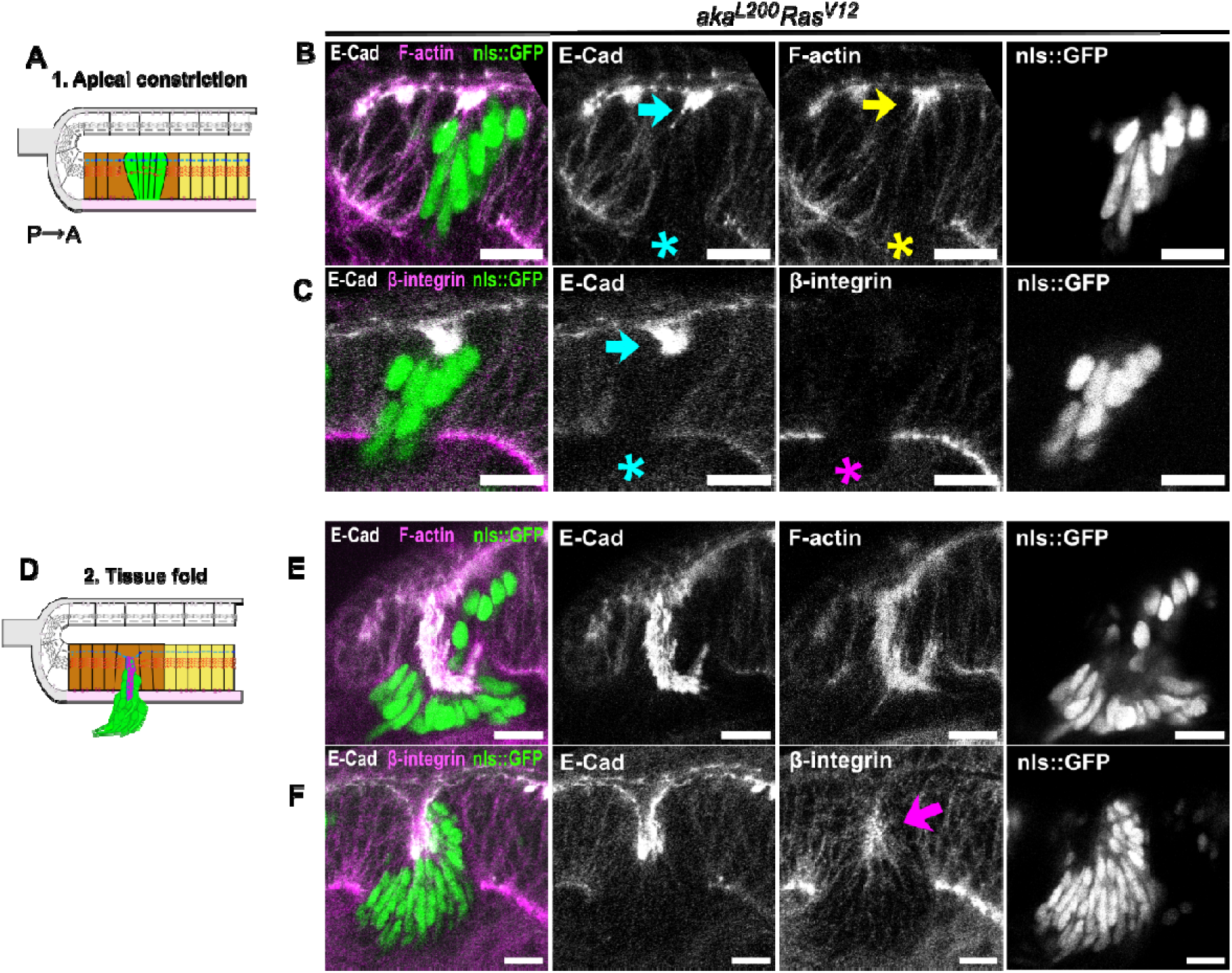
Mutant cells expressing oncogenic Ras form tissue fold anterior to the MF after apical constriction and basolateral delocalization of E-cadherin. (A, D) Schematic of an orthogonal section along the dorso-ventral axis of the *Drosophila* eye disc in the differentiated region of the pseudostratified epithelium (brown), anterior to undifferentiated cells (yellow). Mutant cells (green) apically constrict (A) to then form tissue folds (D). (B, C, E, F) Heterozygous eye disc immunostained with anti-E-cad (grey) and phalloidin (B, E, H magenta) or anti-β-integrin (C, F, magenta). Homozygous mutant cells for aka (*aka^L200^*) and oncogenic Ras cells express the clone marker nls::GFP (green). (B, C) At xx h AEL, mutant cells posterior to the MF appear apically constricted (blue arrows) and have lost the basal loss of E-cad (blue asterisk). Anchoring to the extracellular matrix is lost, as observed by the basal loss of β-integrin (pink asterisk) and actin stress fibers (yellow asterisk). (E, F) Mutant cells continue to exhibit apical constriction and loss of basal adhesion and invaginate. Apical enrichment of β-integrin (pink arrow) is observed, suggested preferential adhesion between mutant cells is present. The scale bars represent 10 μm. All images are single confocal optical sections with posterior to the left, anterior to the right. See also Figure S3.

### Rosettes are composed of differentiated cells delaminating from SJ-deficient *Ras^V12^* tumors in the pseudostratified epithelium

We next studied cell identity and polarity markers in addition to junctional components to decipher the origin of rosettes formed upon basal delamination of SJ-deficient tumors. Rosettes systematically appeared as a cluster of mutant cells organized radially, detached from the cell layer and located basal to the eye disc. We stained for Elav, a marker of early neural differentiation (Robinow & White, 1991), and the atypical protein kinase C (aPKC), a component of the apical Par polarity complex (Wodarz et al., 2000). Anterior to the MF, intermediate-size clones undergoing apical constriction were formed by a homogenous population of undifferentiated Elav-negative mutant cells (Fig. 4A, B). Moreover, basolateral E-cad delocalization at this stage was accompanied by a partial delocalization of aPKC to the basal pole, suggesting a partial loss of polarity (Fig. S3A-B). Similarly, large clones in this region forming tissue folds comprised a homogenous population of undifferentiated, Elav-negative mutant cells (Fig. 4C, D). In these clones, E-cad basolateral delocalization and aPKC partial basal delocalization suggested an overall alteration of the apico-basal polarity (Fig. S3C, D). Conversely, posterior to the MF, small clones undergoing apical constriction and basally depleted in E-cad were composed of a homogenous Elav-positive mutant cell population (Fig. 4E, F). In these clones, aPKC was apically restricted, suggesting polarity is unaltered (Fig. S3E, F). Similarly, large clones forming tissue folds in this region also comprised a homogenously Elav-positive cell population (Fig. 4G, H). In these clones, polarity was preserved and aPKC appeared to be restricted apically like in wild-type neighbors (Fig. S3G, H). Basal delaminated rosettes systematically appeared basal to the differentiated region of the pseudostratified epithelium formed by Elav-positive, polarized cells (Fig. 4I, J). In rosettes, aPKC was restricted to the apical domain, where adhesion components E-cad and β-integrin were also localized (Fig. S3I-K). Based on their location, the expression of Elav and their polarity status, we propose rosettes originate from SJ- deficient tumor located in the pseudostratified epithelium, posterior to the MF, that undergo basal collective delamination.

**Figure 4.**
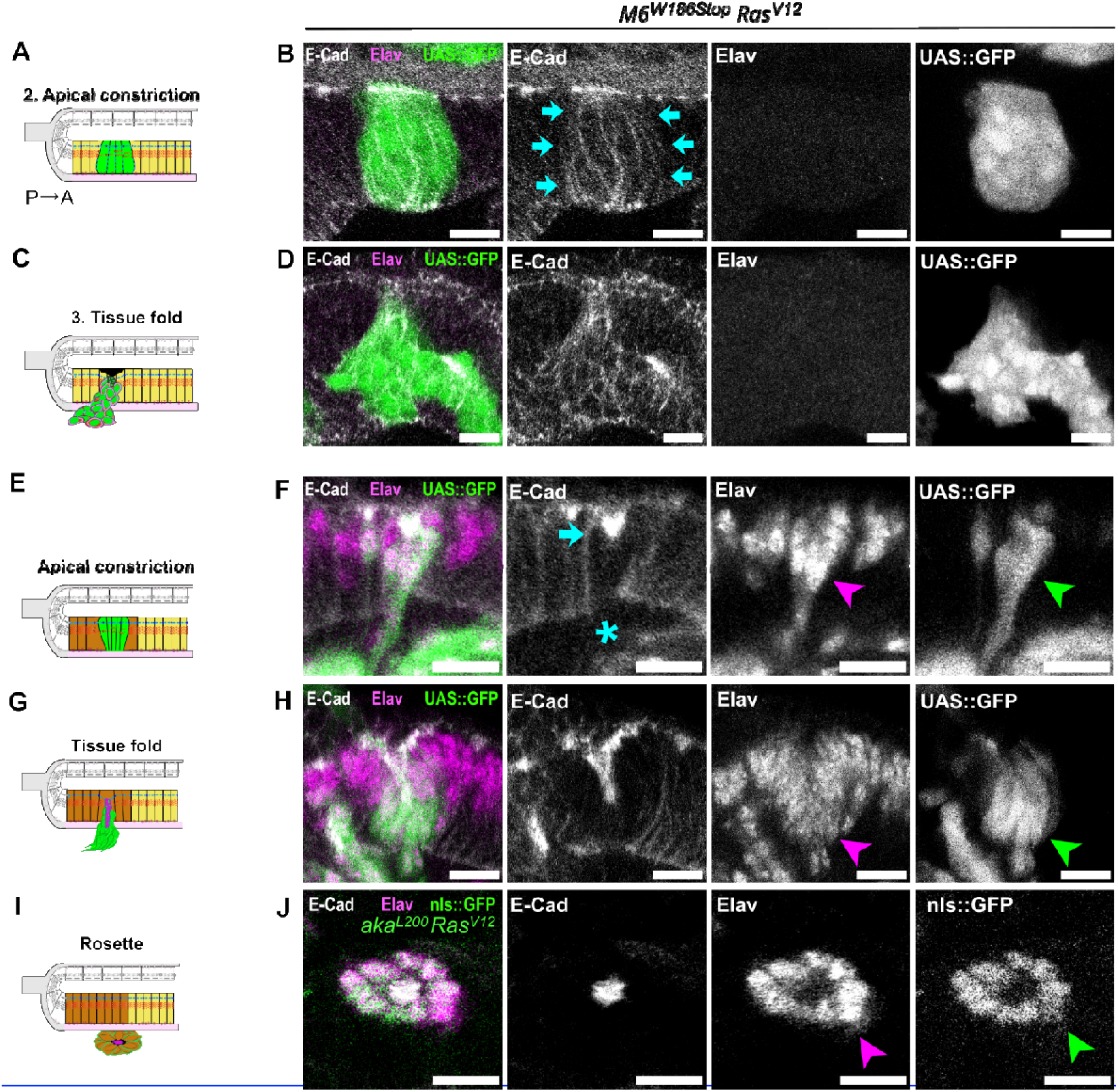
Rosettes are exclusively formed by polarized differentiated cells while tissue folds arise from either undifferentiated unpolarized cells, or differentiated polarized cells. (A, C, E, G, I) Schematic of an orthogonal section along the dorso-ventral axis of the *Drosophila* eye disc of collective mutlicelluar rearrangements with basal directionality that arise from the pseudostratified epithelium (brown, yellow). (B, C, E, F) Heterozygous eye disc immunostained with and stained with anti-*E-cad* (grey) and anti-Elav (pan-neuronal cell marker, magenta). Homozygous mutant cells for M6 (*M6^W186Stop^*) and oncogenic Ras cells express the clone marker UAS::GFP (green). (B) Clones in the undifferentiated region of the pseudostratified epithelium anterior to the MF, when they undergo apical constriction with basolateral spreading of E-cad (blue arrow), are formed by clusters of xx cells, homogenously negative for Elav. (D) Tissue folds formed anterior to the MF are also homogenously negative for Elav. (F) Clones in the differentiated region of the pseudostratified epithelium (green arrowhead), posterior to the MF, are homogenously positive for Elav (magenta arrow head) when they initially undergo apical constrict (blue arrow) and loss of basal adhesion to the extracellular matrix (blue asterisk). (F) Tissue folds formed posterior to the MF (green arrowhead) are formed by homogenously Elav-positive cells (magenta arrowhead). Rosettes are exclusively formed by Elav-expressing cells (magenta arrowheads, green arrowhead). The scale bars represent 10 μm. All images are single confocal optical sections with posterior to the left, anterior to the right. See also Figure S3.

### SJ-deficient *Ras^V12^* tumors delaminate apically from the flat epithelium

Our spatiotemporal analyses on fixed specimen revealed that SJ-deficient tumor generated in the flat epithelium systematically underwent apical collective delamination in the lumen. First, at the beginning of L3, in the early stages of apical collective delamination, *M6^VW186Stop^Ras^V12^* clones had an average size of 6 cells (± 2), underwent apical constriction and acquired a dome-like shape on the basal side. On the apical side, clones formed a neck in the plane of adherens junctions and emitted cytoplasmic apical protrusions into the lumen. Because of their morphology, we have named them octopus-shaped clones. This stage is characterized by the lateral enrichment of E-cadherin and β-integrin between *M6^VW186Stop^Ras^V12^* cells (Fig. 5A-C). Next, until 115hAEL, clone sized increased to 13 cells (± 5) and mutant cells were observed to translocate into the lumen through the apical neck (blue arrow, Fig. 5E). Mutant cells remained all connected, indicative of a collective delamination, and β-integrin appeared enriched in the entire clone (Fig. 5D-F). More precisely, β-integrin was enriched at the cell membrane of mutant cells in the flat epithelium, in those delaminating through the apical neck and in those successfully deviated in the lumen. From 114hAEL until 120hAEL, in the last step of apical collective delamination, cluster size was 18 cells (± 5) and mutant cells were totally detached from the cell layer (Fig. 5G, H). Mutant cells appeared collectively delaminated in the lumen and enriched in adhesion proteins like β-integrin (Fig. 5I). Altogether, these *in vivo* results show SJ-deficient tumors exclusively undergo basal collective delamination when originated in the pseudostratified epithelium and apical collective delamination when formed in the flat epithelium. This suggest that cell morphology and the sequential remodeling of cell junctions are key to ensuring the cohesion and directionality of collective delamination of SJ-deficient tumors alive out of an epithelium.

**Figure 5.**
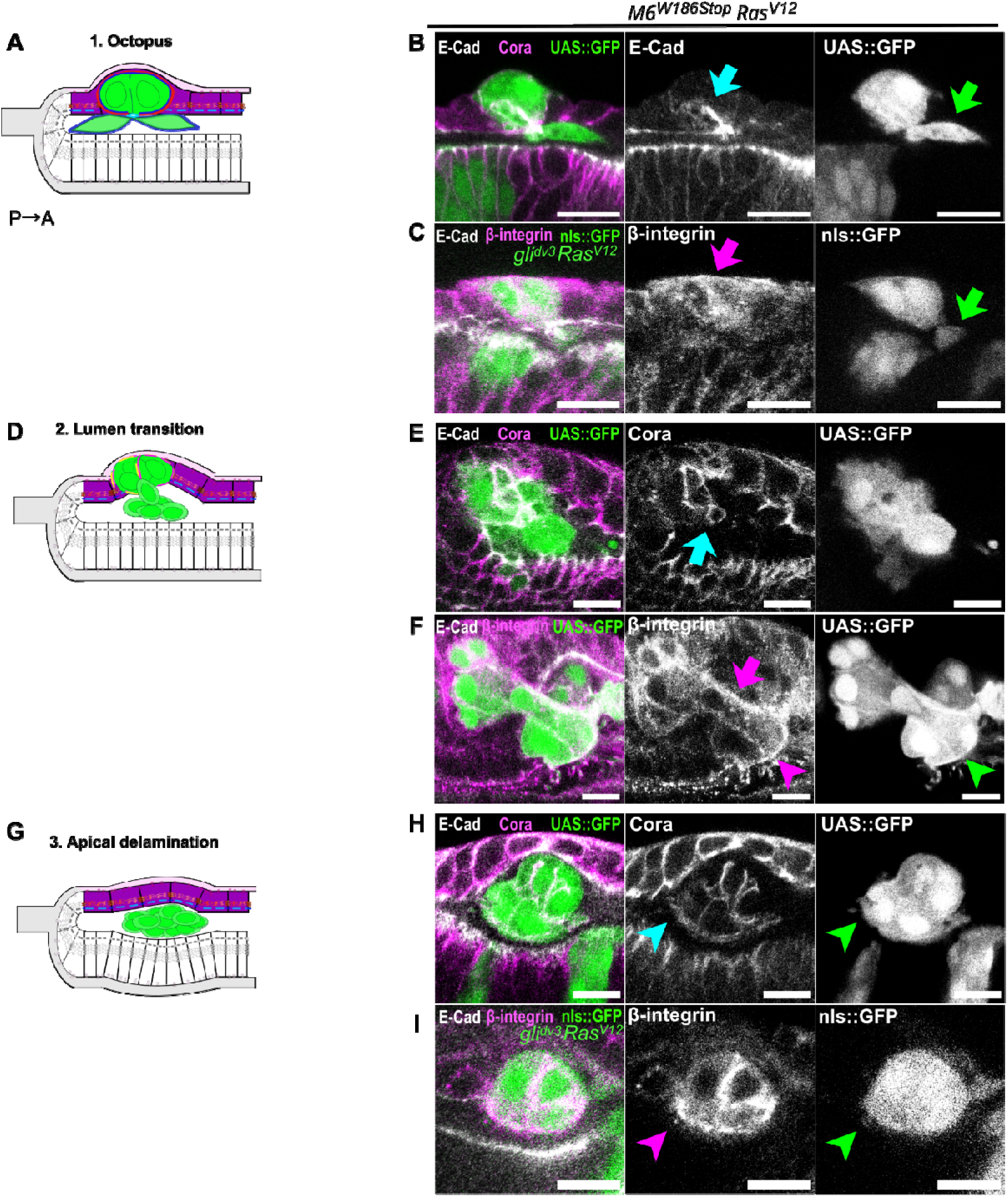
Apical Delamination of mutant cells from the flat epithelium form an octopus and an apical neck before translocating in the lumen. (A, D, G) Schematic of an orthogonal section along the dorso-ventral axis of the *Drosophila* eye disc of the flat epithelium (purple). Mutant cells (green) apically constrict (A), progressively translocate into the lumen (D) and ultimately appear completely detached in the lumen (G). (B, C, E, F, H, I) Heterozygous eye disc immunostained with anti-E-cad (grey) and anti-Cora (B, E, H, magenta) or anti-β-integrin (C, F, I, magenta). Homozygous mutant cells for M6 (*M6^W186Stop^*) and oncogenic Ras cells express the clone marker UAS::GFP (green). (B, C) During the time window 89-99hAEL, mutant cells in the flat epithelium (green) appear apically constricted and acquire a dome-shape, termed the octopus, which exhibits basolateral enrichment of E-cad (green arrow) and β-integrin (magenta arrow), and apical lumen protrusions (green arrow). (E, F) Between 90-115hAEL, mutant cells progressively translocate into the lumen (green arrowhead) through a ring-like structure enriched in Cora (blue arrow). β-integrin enrichment is present all throughout the clone, in cells moving from the flat epithelium into the lumen (pink arrow) and cells located in the lumen (pink arrowhead) as (H, I) Lastly, between 114-120hAEL, mutant cells (green) appear complelty detached in the lumen, with no conneciotn to the flat epithelium (blue arrowhead) and enriched in β- integrin (pink arrowhead). The scale bars represent 10 μm. All images are single confocal optical sections with posterior to the left, anterior to the right.

### Computational modeling recapitulates how flat and pseudostratified epithelia respond differently to epithelial destabilization events

To further assess the putative importance of the tissue organization and of the various events taking place prior to cell delamination, we turned to computational modelling. We adjusted a recently developed 2D agent-based model of epithelial dynamics (Despin-Guitard et al., 2024; Plunder et al., 2024) (Fig. 6A, see Methods, Supplementary Table 1) to obtain a flat monolayer (Fig. 6B, movieS1) or a short pseudostratified organization (Fig. 6C, movieS1) as observed in the eye-antennal imaginal discs. We implemented for the clone cells apical constriction (event ***ApC***), loss of apical cell-cell adhesion (event ***A***) and loss of basal adhesion to the cell-matrix (event ***B***) and monitored the rate of apical and basal delamination (Fig. 6D). We started by simulating these events either independently or occurring sequentially in various orders, in a single cell, in both types of epithelia (Fig. 6E). All events are progressively implemented over 20 hours of simulation and the epithelia are left to further develop for an extra 30h after the last event took place. ***ApC*** alone does not trigger delamination on either side but leads to a basal displacement of cell nuclei in the pseudostratified tissue (Fig S4). Interestingly, overall epithelial destabilization in the flat epithelium preferentially leads to apical delamination (Fig. 6E), unless ***B*** is the first event (Fig. 6E) in which case delamination in randomized. By contrast, in the short pseudostratified tissue basal delamination is systematically the main outcome regardless of the scenario (Fig. 6E-F).

**Figure 6.**
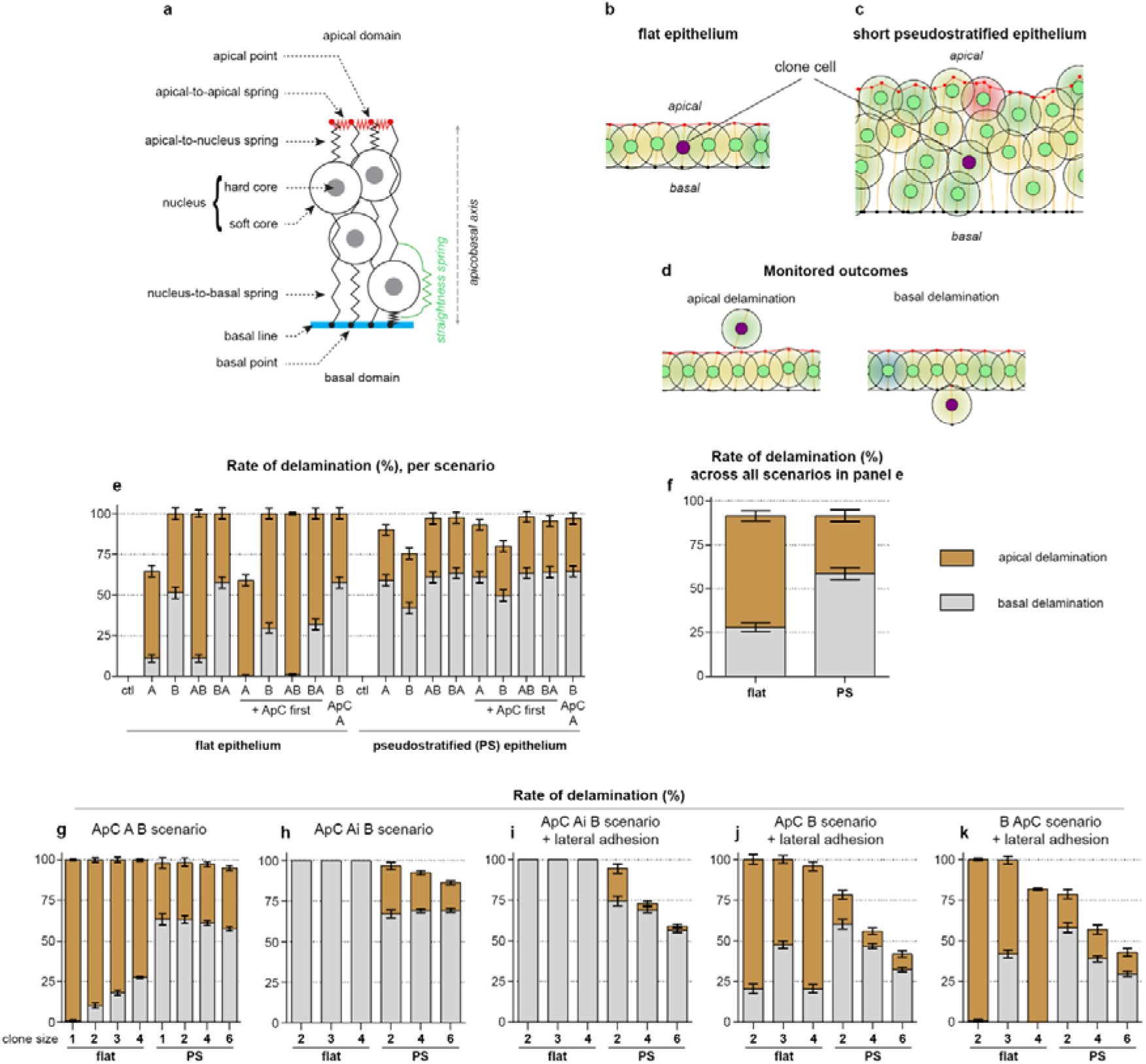
Differential sensitivity of flat and pseudostratified tissues to simulated epithelial destabilization. (A) Diagram representing the simulated cells and the various adjustable springs control their viscoelastic properties. (B, C) Examples of modelling outputs showing the epithelium in flat or short pseudostratified configuration. Control cells have green nuclei, clone cells have magenta nuclei. (D) Examples of apical and basal delamination upon epithelia destabilization. (E) Mean rates of delamination with standard error of the mean for each tested scenario in single cells. (F) Global rate of delamination for all simulations shown in panel E. (G-K) Mean rates of delamination with standard error of the mean for all indicated scenarios tested in clones of various sizes. Each simulation was performed 200 times. A, loss of apical adhesion between clone cells and between clone cells and control cells; Ai, loss of apical adhesion only at the interface between clone cells and control cells; ApC, apical constriction; B, loss of basal adhesion; PS, pseudostratified.

### Clone size and lateral adhesion contribute to decision making in collective cell delamination

Next, we asked whether the outcome of epithelial destabilization was sensitive to clone size. In 3D, clone size is tripled to the corresponding size in the 2D-agent model. Hence, we utilized our *in vivo* data to restrain the size of cell populations undergoing delamination *in silico* to study biologically relevant scenarios. We first focused on the ***ApC A B*** scenario, which involves all events likely to happen *in vivo* based on the above characterized spatiotemporal analyses and leads to the expected directionality of delamination in each tissue, respectively apical and basal for the flat and pseudostratified epithelia. Increasing the clone size in the flat epithelium progressively increases the rate of basal delamination up to 25% of the cells (Fig. 6G, movieS2). By contrast, the impact of increasing the clone size in the pseudostratified epithelium was marginal (Fig. 6G, movieS2). One striking observation from these simulations is that there is a splitting of the clones, i.e. not all cells from the clone extrude on the same side of the epithelium (MovieS2). This was not observed *in vivo*. This strongly suggests that *in vivo* collective cell delamination requires cell-cell adhesion between mutant cells to be maintained.

### Lateral adhesion between delaminating mutant cells and their wild-type neighbors ensures directionality

To assess the impact of cell-cell adhesion, we introduced a new event called ***Ai*** (for event ***A*** only at the interface between control cells and clone cells). During this event, clone cells lose apical cell-cell adhesions only at the interface between them and the control cells of the tissue. Normal apical cell-cell adhesions between clone cells are maintained. When ***Ai*** occurs, the two control cells on either side of the clone form a new bond and “heal” the local wound (see Methods). In the pseudostratified tissue, maintaining adhesion between clone cells mildly reduced the rate of apical delamination but was not sufficient to prevent clone splitting (Fig. 6H, movieS3). However, in the flat epithelium, maintaining cell-cell adhesion between clone cells, together with ***Ai*** occurring, led to a complete reversal of the directionality of delamination (Fig. 6H, movieS3). To further implement adhesion between clone cells, we added a maximum distance rule between clone cells such that upon detachment from the matrix (***B***) clone cells are prevented from drifting away from each other, mimicking maintenance of a strong lateral adhesion. This did not affect the outcome of the simulations in the flat epithelium (Fig. 6I, movie S4). By contrast, in the pseudostratified tissue, enforcing lateral adhesion suppressed clone splitting and made apical delamination a rare case (Fig. 6I, movieS4).

*In vivo*, delamination occurs while E-cad is still detected in clone cells suggesting that adhesion between clone cells and control cells may not systematically be lost. Therefore, we also tested alternative scenarios in which only events ***ApC*** and ***B*** occur (Fig. 6 JK). In the flat tissue, the maintenance of adhesion between mutant and wild-type cells made apical delamination the main outcome. In the pseudostratified epithelium, the overall delamination rate is progressively reduced as clone size increases and the apical delamination, which was a rare event in the ***ApC Ai B*** scenario, now represents a third of all outcomes.

### Basal membrane rigidity regulates directionality of delamination exclusively in the flat epithelium

*In vivo*, all events seem to occur in a relatively short time window (circa 3-4h). Therefore, we decided to implement various scenarios in the pseudostratified epithelium allowing events to take place within a 4h time frame (Fig. S5 A-D). Rushing cells to perform the various events did not affect the overall outcome of each tested scenario. Next, we tested an extreme case with ***A*** and ***B*** occurring simultaneously in each epithelium (Fig. S5E). Again, in the pseudostratified epithelium the effect is marginal as basal delamination dominates. However, in the flat tissue having ***A*** and ***B*** implemented simultaneously randomizes delamination in single cells and slightly favors basal delamination in larger clones (Fig. S5E).

These simulations indicate that epithelial destabilization in the short pseudostratified tissue primarily leads to basal delamination with marginal effects from the order and timing of events. They also show that extensive cell-cell adhesion (apical and lateral) is required for delamination to occur collectively. By contrast, in the flat epithelium, epithelial destabilization in a single cell preferentially leads to apical delamination but the tissue is highly sensitive to the type of events taking place, in which order they occur and to clone size. For instance, modulating adhesion between clone cells or between clone cells and control cells was sufficient to toggle between apical and basal delamination.

So far, all simulations were run in a permissive context in which the basal compartment does not act as a physical barrier. However, *in vivo*, both flat and pseudostratified layers of the eye-antennal disc are anchored to an extracellular matrix. To test the impact of such environment, we assessed the outcomes of some of the previously tested scenarios in the context of a non-permissive basal matrix acting as a physical obstacle and as a substrate, forcing cells to experience a small degree of confinement (Fig. 7A-C, movieS5). As expected the non-permissive matrix prevents basal delamination. Interestingly, in the flat tissue, this was sufficient to promote apical delamination for all scenario including the ones that have an intrinsic bias towards basal delamination (Fig. 7A, compared to Fig. 6I). However, in the pseudostratified epithelium preventing basal delamination is not sufficient to drive apical delamination (Fig. 7A-C). To assess whether the non-permissive matrix nonetheless influenced the position of clone cells in the pseudostratified tissue, we monitored the rate of basal positioning which indicates the percentage of cells with their nuclei located in a more basal position than the average position of the control cells (Fig. 7D, E). Interestingly, all scenarios tested in the pseudostratified epithelium led to basal positioning of cells (Fig. 7E). This indicates that the intrinsic basal directionality induced by epithelial destabilization observed in the pseudostratified tissue still occurs despite actual delamination being prevented by the non-permissive matrix. Overall, in the flat epithelium there are various possible outcomes of epithelial destabilization depending on the type and order of events but these are likely to be cancelled by the presence of the matrix acting as a physical barrier which strongly biases delamination towards the apical side. By contrast, all scenarios tested in the pseudostratified tissue led to basal accumulation of cells that either delaminate if the matrix is permissive or remain in the tissue is the matrix acts as a physical barrier.

**Figure 7.**
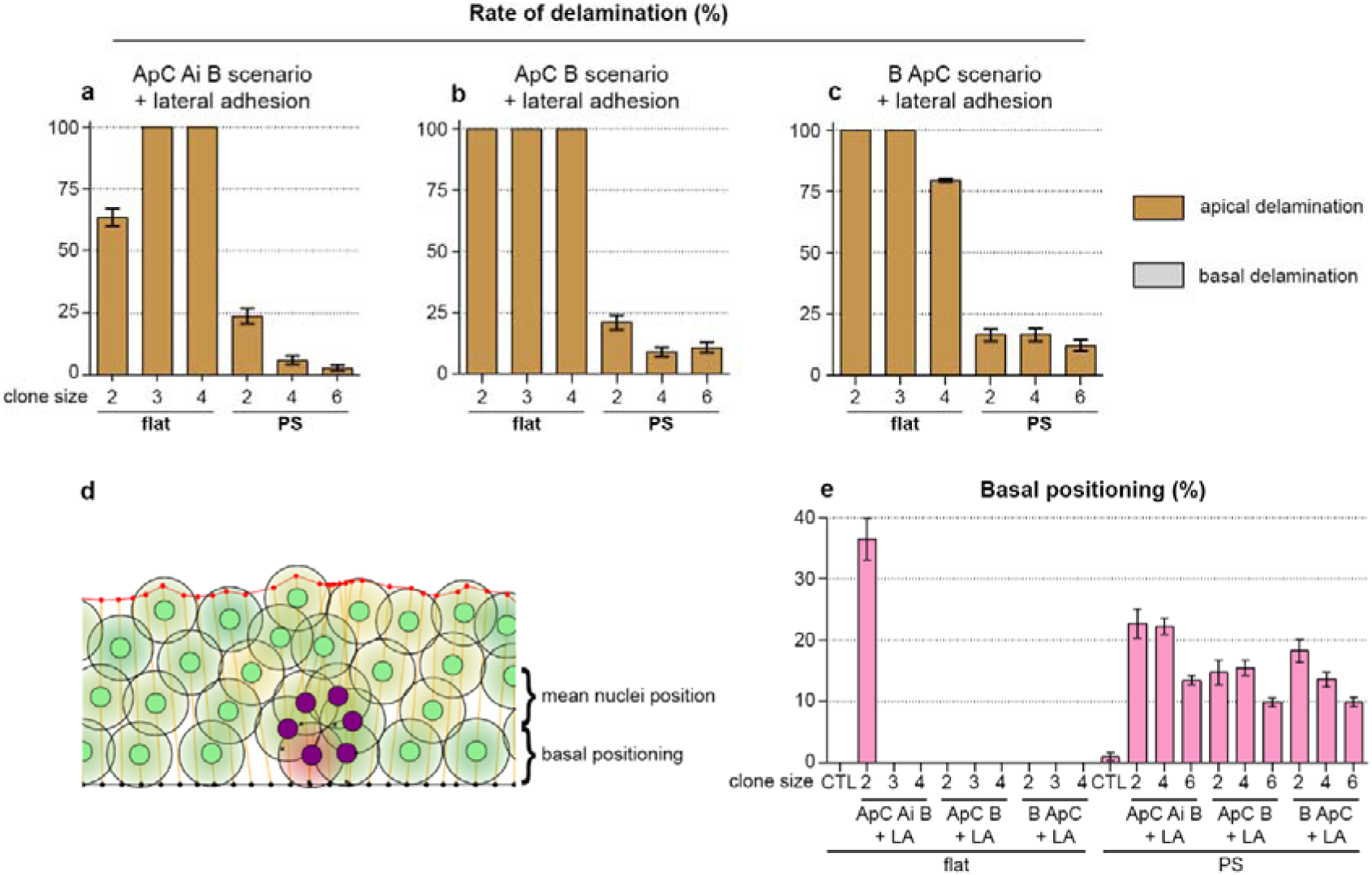
A non-permissive basal compartment converts epithelial destabilization in the flat tissue into apical delamination. (A-C) Mean rates of delamination with standard error of the mean for all indicated scenarios tested in clones of various sizes in the presence of a non-permissive basal compartment. Each simulation was performed 200 times. (D) Example of a clone (magenta) that underwent the ApC B and lateral adhesion scenario, note that only some of the clone cells are considered basally positioned. (E) Rates of basal positioning for each tested scenario. A, loss of apical adhesion between clone cells and between clone cells and control cells; Ai, loss of apical adhesion only at the interface between clone cells and control cells; ApC, apical constriction; B, loss of basal adhesion; LA, lateral adhesion; PS, pseudostratified.

### Mechanical forces and preferential adhesion are at interplay during apical collective delamination of tumors

Mathematical modelling of epithelial destabilization in a flat tissue revealed loss of adhesion between mutant cells and wild-type neighbors is sufficient to revert the directionality of apical delamination despite mutant cells showcasing preferential adhesion. Counterintuitively, this suggests that intraconal adhesion between mutant cells and intercellular adhesion with wild-type neighbors are required to elicit apical collective delamination out of a flat epithelium. *In silico* results coincided with *in vivo* observations and noted apical constriction to be an important feature during collective delamination. To gain more insight on the underlying mechanisms of apical collective delamination, we studied the activation of the actomyosin cytoskeleton network, using phospho-Myo as a proxy, together with junctional markers. As aforementioned, during early apical collective delamination, clones underwent apical constriction and clustered in a dome-like shape (Fig. 8A). Mutant cells were embedded in the cell layer and emited apical lumen protrusion, hence the term “octopus” to describe this stage. Phospho-Myosin was enriched at the periphery of the octopus, at the interface between mutant clones and their wild-type counterparts (Fig. 8B, yellow arrows). We also observed cytoskeleton remodeling at the tip of lumen protrusions (Fig. 8B yellow and green arrowheads), suggesting both non-cell autonomous and cell-autonomous forces were at play early during apical collective delamination events. During metamorphosis of the pupal abdomen, an extracellular ring of actomyosin forms non-cell autonomously to extrude larval epidermal cells (Teng et al., 2017). This supracellular structure has also been described as part of the tissue clearance mechanisms used to eliminate the epidermal cells of the pupal notum, which eventually extrude and die by apoptosis (Villars et al., 2022). However, no caspase activation was detected at the octopus stage (Fig. 8C), suggesting apical collective delamination occurs independently of caspase-activation and involved cytoskeleton remodeling at the periphery of mutant cells. Mathematical modelling revealed that lateral adhesion between mutant and wild-type cells was essential for collective apical delamination in a flat epithelium. At later stages of apical collective delamination, when cells collectively translocated in the lumen, we observed *in vivo* a ring-like structure enriched in adhesion proteins like Coracle and E-cad (Fig. 8D, E, blue arrows). We termed this structure the neck, through which cells left the flat epithelium and delaminated, translocating their nucleus and cytoplasm into the lumen (Fig. 8D, E, blue arrowhead). Three-dimensional reconstruction and quantification revealed the neck to be on average 3µm in diameter (Fig. 8F). This value is in range with *in vivo* studies cell migration of dendritic cells in the mouse ear skin, for which a minimal diameter of 2µm suffices (Raab et al., 2016). Altogether, our *in silico* and *in vivo* results suggest tumors with disrupted SJ integrity undergo basal collective delamination when originated in a pseudostratified epithelium regardless of the order of junctional remodeling. Conversely, SJ-deficient tumors formed in a flat epithelium exclusively undergo apical collective delamination. Moreover, the directionality of collective delamination in flat epithelia is highly to the sequence in which junctional remodeling is orchestrated and preferential adhesion between mutant cells, and to their wild-type neighbors ensures collectiveness and completion of the process.

**Figure 8.**
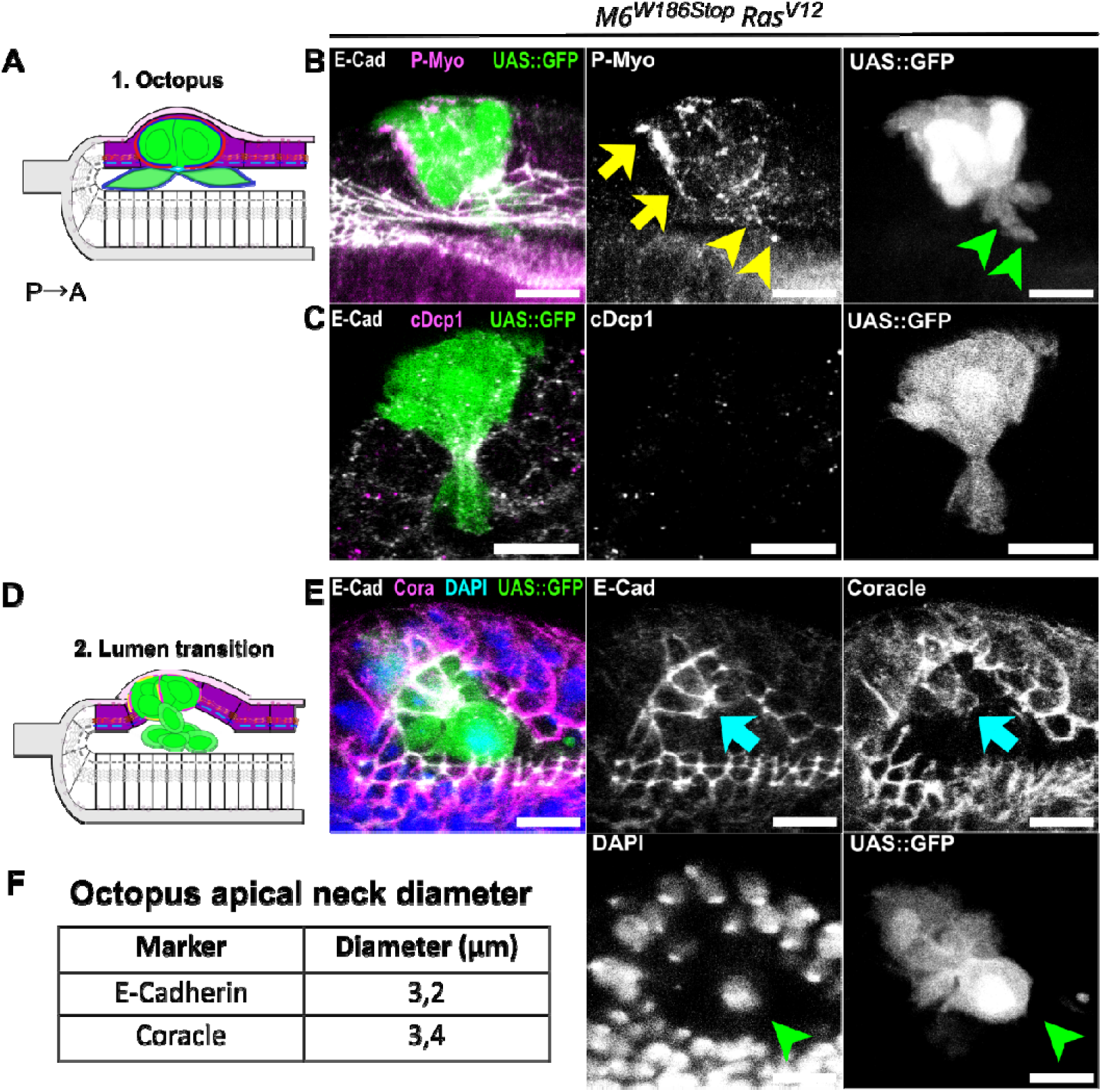
Distribution of actomyosin force generator and preferential adhesion guide apical cell delamination. (A, D) Schematic of an orthogonal section along the dorso-ventral axis of the *Drosophila* eye disc of the flat epithelium (purple). Mutant cells (green) apically constrict (A) into a dome-shaped octopus with apical protrusions and progressively translocate into the lumen (D). (B, C, E) Heterozygous eye disc immunostained with anti-E-cad (grey) and anti-P-Myo (B, magenta) or anti-Drosophila caspase-1 (DCP-1) (C, magenta). Homozygous mutant cells for M6 (*M6^W186Stop^*) and oncogenic Ras cells express the clone marker UAS::GFP (green). (B, C) During apical collective delamination, at the octopus stage, P-Myo appears enriched at the interface between mutant cells and their wild-type counterparts (yellow arrows) as well as at the tip of lumen protrusions (yellow arrowheads). No signs of caspase activation at this stage are detected, suggesting apical collective delamination occurs independently of apoptosis activation. (E) During translocation of cells form the flat epithelium into the lumen a ring-like structure enriched in adhesion proteins (blue arrows) termed the “neck” it formed, with average diameter of 3µm for cells to delaminate in the lumen (green arrowheads). (F) Quantification of neck size performed on eye-antennal imaginal disc stained with anti-E-cadherin and anti-Coracle. The scale bars represent 10 μm. All images are single confocal optical sections with posterior to the left, anterior to the right.

## Discussion

### SJ loss promotes cell elimination in the larval eye disc

Here, using an *in vivo* cooperative tumorigenesis system in the *Drosophila* larval eye disc, we found that loss of SJ integrity alone can trigger caspase-induced elimination during L2 and L3 stages. This elimination takes place before the developmentally programmed cell-death occurs during pupal metamorphosis (Wolff & Ready, 1991). Hence, SJ-deficient clones are removed from the eye independently of this process. SJ play a major role in the establishment of the blood-retinal-barrier, sealing light-sensitive photoreceptors from the circulating hemolymph to enable phototransduction (Banerjee et al., 2008). Our results reveal single mutant cells deficient for bi- or tri-cellular SJ are prone to die by apoptosis in the flat and pseudostratified epithelium. We reasoned this could be due to cell competition mechanisms, known to preserve tissue homeostasis by removing potentially hazardous cells from the cell layer (Baker, 2020; Levayer, 2018; Morata, 2021). Loss of SJ integrity could decrease the relative fitness of mutant cells in comparison to their adjacent wild-type neighbours due to changes in the molecular fitness fingerprints or mechanical properties. This would in turn trigger cell competition and promote the elimination of SJ mutant cells via caspase-activation. At the molecular level, loss of fitness in mutant cells could be due to the mislocalization of SJ components. In the pupal notum, disruption of bicellular SJ upon loss of *nrv2* leads to a spreading of Gli from tricellular SJ into the bicellular SJ domain (Esmangart de Bournonville & Le Borgne, 2020). In the wing disc which, Gli mislocalization promotes Dlg downregulation and ultimately triggers cell elimination (Padash-Barmchi et al., 2010, 2013). Alternatively, loss of SJ integrity in the eye disc could be accompanied by the dysregulation of the ESCRT/retromer system, as described in the pupal notum (Esmangart de Bournonville et al., 2024). This would result in the accumulation of ubiquitinylated substrates and proteotoxic stress, triggering cell competition as observed in the wing disc (Baumgartner et al., 2021). Furthermore, differences in the rate of proliferation between two cell populations in a mosaic tissue triggers cell competition and elimination of the slower cycling population (Morata & Ripoll, 1975). Cora was first identified as a dominant negative suppressor of the Epidermal Growth Factor Receptor (EGFR) signaling pathway responsible for promoting cell survival (Fehon et al., 1994). *cora* loss-of-function decreases cell proliferation, triggering cell elimination in the eye-antenna and wing imaginal discs without causing major tissue disruption (Ward et al., 2001).

### Cooperative SJ loss and oncogenic Ras promote collective delamination of living

Strikingly, our findings show that upon loss of SJ in cooperation with Ras malignant transformation, SJ-deficient tumors do not die. Instead SJ-deficient tumors undergo different types of collective cell movements, namely tissue folding and apical or basal collective delamination of living cells. This is, to our knowledge, one of the first descriptions of a permissive function for SJ together with oncogenic Ras in the initiation of cell delamination. We reasoned Ras overactivation in SJ-depleted cells bestows a fitness advantage and masks the “loser” fitness status. This would in turn transform SJ-mutant *Ras^V12^* cells in “super-competitors”, a phenomenon observed in the context of tumor progression (Cong & Cagan, 2024). Ras overactivation in the wing disc does not generate significant overgrowth but instead upregulates the oncogene dMyc, rendering cells refractory to apoptosis due to the ectopic trans-activation of *head involution defective (hid)* by the Ras-MAPK pathway (De La Cova et al., 2004; Moreno & Basler, 2004; Pinal et al., 2018). Ras overactivation in the eye disc also downregulates *hid,* promoting cell survival (Bergmann et al., 1998) while *Ras^V12^*transformed cells trigger compaction and elimination wild-type neighbors in the pupal notum (Levayer et al., 2016; Moreno et al., 2019).

### Folding morphogenesis exhibits preferential cell adhesion mediated by E-cadherin and β-integrin

We performed spatiotemporal characterization and dissected the cellular steps of collective cell movements elicited by SJ-deficient *Ras^V12^*tumors. Our results show tissue folds exclusively form in the pseudostratified epithelium of the eye disc. This tissue differentiates with the posterior-to-anterior MF progression and is composed of two cell populations: undifferentiated progenitors, anterior to the MF, and differentiated cells, posterior to the MF (Warren & Kumar, 2023). In this epithelium, folding morphogenesis of SJ-deficient *Ras^V12^* tumors comprises junctional remodeling at apical, lateral and basal domains. This mechanical coupling has also been observed in other tissues (Denk-Lobnig & Martin, 2020; Tozluoglu & Mao, 2020). We describe how junctional rearrangements differ depending on cell identity and describe the presence of β-integrin and E-cad-mediated adhesion during tissue folding. Firstly, Elav-negative SJ-depleted *Ras^V12^* tumors anterior to the MF exhibit a partial loss of polarity and undergo apical constriction. Concomitantly the lateral cortex is remodeled, evidenced by E-cad spreading and P-Myo enrichment, while focal adhesions preserve basal anchoring to the extracellular matrix. Basolateral enrichment of the phosphorylated regulatory light chain of non-muscle type II myosin (P-Myo) suggests an increase in contractility of the actomyosin cytoskeleton at this cortical domain (Fox & Peifer, 2007; Mao et al., 2013). The partial loss of polarity and junctional remodeling events persist, and apical constriction continues. Moreover, tissue folds appear enriched in the focal adhesion component β-integrin at apical and lateral domains. This suggests the presence of preferential adhesion mediated by E-cad and β-integrin. In *Xenopus,* during the migration of neural crest cells, cadherins have been found to function at focal adhesion and mediate adhesion to the extracellular matrix (Langhe et al., 2016). It would be interesting to further explore whether integrins can also reciprocate and function to mediate intercellular adhesion. E-cad homotypic interaction is well studied (Brasch et al., 2012) and could be reinforced by the not so well understood homo-oligomerization of β-integrin α-chains, a thermodynamically favorable process previously described in MDCK cells (Manninen, 2015; Maurer et al., 2002; Mehrbod & Mofrad, 2013). In striking contrast, clones located posterior to the MF do not showcase E-cadherin lateral localization at any point during the sequence of junctional remodeling events. SJ-depleted *Ras^V12^* tumors posterior to the MF are Elav-positive and appear well polarized. Tissue folding in these clones starts with the constriction of the apical domain and simultaneous loss of basal anchoring to the extracellular matrix. As apical constriction progresses and the epithelium invaginates, basal loss of β-integrin is concomitant to the specific enrichment or relocalization of β-integrin at the apical domain of mutant cells. This reinforces the notion that strengthening cell-cell adhesion via interplay of E-cad and β-integrin-mediated adhesion may be essential for tissue folding. We cannot rule out the possibility that extracellular matrix may be present at this location, in which case E- Cad and β-integrin could cooperate to provide adhesion, as seen in the *Xenopus* neural crest (Langhe et al., 2016). However, interplay between E-Cad and β-integrin adhesion modules independently of extracellular matrix has been observed *in vitro* during mitosis in cultured cells (Huber et al., 2023). This interplay has also been reported *in vivo*, during embryonic dorsal closure (Narasimha & Brown, 2004) and basal tissue folding of the pseudostratified epithelium in the wing disc (Valencia-Expósito et al., n.d.). Our results further highlight the complexity of tissue folding in pseudostratified epithelia and reveal striking differences of β- integrin and E-cad remodeling concomitant to cell identity and polarity status. The small GTPase Rap1 regulates Cadherin- and Integrin-based adhesion modules during morphogenesis (Boettner & Van Aelst, 2009). During mesoderm invagination in the embryo, Rap1 acts together with Canoe, an adherens junctions-cytoskeleton linker, and regulates β- integrin and E-cad localization (McMahon et al., 2010; Sawyer et al., 2009; Vale et al., 2023). In the eye disc, posterior to the MF, Rap1 affects Notch and EGFR signaling. Rap1 acts through Canoe to regulate E-cad and Baz levels in ommatidia, controlling cell specification and retinal morphogenesis (Gaengel & Mlodzik, 2003; O’Keefe et al., 2009; Walther et al., 2018; Yost et al., 2023). Anterior to the MF, Wingless (Wg) controls the speed of MF progression, which also depends on the expression of integrins by Hedgehog (Hh) and Decapentaplegic (Dpp) signaling. In the wing disc, wg governs tissue folding by basally stabilizing integrins. Hence, the molecular mechanisms regulating folding morphogenesis may be subjected to tissue-intrinsic properties and developmental programs.

### Collective delamination of living cells requires preferential adhesion mediated by E-cadherin and β-integrin

Furthermore, SJ-deficient *Ras^V12^* cells delaminate collectively and alive out of an epithelium, either apically or basally. We have dissected the cellular steps and regulators of the collective delamination of living cells. Our experimental and theoretical results highlight the choice of directionality during delamination is not random, but is instead governed by tissue architecture and preferential cell adhesion. We put forward a detailed molecular plan of junctional remodeling taking place during collective delamination of live tumors. Basal collective delamination of living cells occurs exclusively in the pseudostratified epithelium regardless of the sequence in which junctional complexes are remodeled. This results in the formation of Elav-positive polarized rosettes, radially organized and apically enriched in E- cad, F-actin and β-integrin. We find striking similarities in terms of cell identity, cluster size and remodeling of junctions and the cytoskeleton between rosettes and apically constricted SJ-deficient tumors posterior to the MF. Live imaging will be decisive to unambiguously determine whether rosettes originate indeed posterior to the MF. Conversely, SJ-deficient tumors in the flat epithelium solely undergo apical collective delamination alive out of the epithelium and the sequence of events greatly impacts the directionality of delamination. Our results confirm the synergy between M6 depletion and *Ras^V12^* in promoting cell delamination reported on a previous study (Dunn et al., 2018). Dunn and colleagues identified JNK signaling and Canoe induction of cell-autonomous MyoII contractility via Rho1 activation as mechanical regulators of apical collective delamination (Dunn et al., 2018). The aforementioned study did not include a detailed characterization for the epithelial origin of cells undergoing apical collective delamination. However, they reported clones in the pseudostratified epithelium deviate apically, a phenotype never observed in the experimental set up and analyses conducted in the present study. We performed a detailed spatiotemporal characterization of apical and basal collective delamination and studied markers of junctional and cytoskeletal remodeling, cell identity and polarity status. Larvae were staged from L2 until L3 with a 2h-time window that overlapped across replicates, covering a maximum time of larval development from 76h to 120h. The reported duration of apical collective delamination in (Dunn et al., 2018) was of 190h minutes. Hence, it is unlikely any event of apical or basal collective delamination to have been underscored in our analyses. We quantified nuclear size in clones from the flat and pseudostratified epithelium and compared it to the nuclear size of apically deviated clones. Nuclear size of clones having undergone apical deviation is reminiscent to that of clones from the flat epithelium, which is significantly bigger than clones in the pseudostratified layer. Moreover, we performed mathematical modeling using a powerful EMT-simulator adapted to the eye disc architecture. Both *in vivo* and *in silico* results reveal apical collective delamination events exclusively originate from the flat epithelium, and never from the pseudostratified one. The reason for the discrepancy between the origin of apically delaminating clones between the aforementioned study (Dunn et al., 2018) and our results remains unknown. Live imaging with clonal and junctional markers would help clarify this issue. In any case, our studies expand on the role of bi- and tricellular SJ components and we report a non-occluding function of SJ in preventing cell extrusion in the eye disc. Loss of SJ integrity synergizes with oncogenic Ras and promotes collective delamination of tumors alive out of the epithelium.

Collective apical delamination in the flat epithelium starts with SJ-deficient *Ras^V12^* tumors acquiring a dome-like shape and forming an “octopus”. Basolateral preferential adhesion between mutant cells is mediated by E-cad and β-integrin. At the apical domain, cell protrusions are emitted in the lumen. Concomitantly, an apical neck is formed and contains Cora and E-cad. Mutant cells delaminate collectively and alive out of the flat tissue through this neck, deviating in the lumen while remaining cohesive. The presence of P-Myo and phalloidin at the tip of apical lumen protrusions suggests the SJ-deficient tumor could exert pulling forces at the end of cell protrusions. In addition, P-Myo is enriched at the lateral pole, all along the rim of the SJ-deficient tumor, forming what looks like a supracellular cable. The contraction of such supracellular cable could exert pushing forces that would in turn contribute to apical delamination. This supracellular cable could be formed by wild-type neighbors, as observed during cell extrusion in the pupal abdomen and notum (Levayer et al., 2016; Teng et al., 2017). Importantly, *in silico* mathematical modelling of the flat epithelium highlights cell adhesion between wild-type and mutant cells to be essential for apical collective delamination. This suggests E-cad mediated apical adhesion observed *in vivo* at the neck is key for this process. The relative contribution of mechanical forces in this process awaits further studies. Yet, cell-autonomous pulling forces could be exerted at the tip of apical lumen protrusions concurrent to non-cell autonomous pushing forces located at the mutant-wild-type interface of the actomyosin supracellular cable. Our study hence suggests mechanical tension exerted by neighbors may not be restricted to apoptotic-dependent cell extrusion (Villars & Levayer, 2022). Therefore, non-cell autonomous forces may be required for apical collective delamination of living cells. It would be very informative to prevent force activation cell-autonomously in mutant cells or non-cell-autonomously in wild-type neighbors to further elucidate their contribution to this process. Moreover, studying actomyosin force regulators such as α-catenin, fluorescently tagged and expressed exclusively in the mutant or wild-type cells, would clarify the source of mechanical activation and force generation. This would clarify whether supracellular myosin cables form exclusively during caspase-dependent cell extrusion or whether this mechanism is overridden by “super-competitor” tumors and required for living cells to delaminate collectively. Tumor delamination is the first step in metastasis onset. Further characterization of the molecular regulators involved may shed light on ubiquitous mechanisms required for collective cell rearrangements and co-opted by cancer cells to drive tumor malignancy. Moreover, the direction in which cells leave an epithelium can be critical for determining cell fate (Gu & Rosenblatt, 2012). TJ in vertebrates, the somewhat analogous to invertebrates’ SJ, are located apical to adherens junctions. Oncogenic transformation triggers basal extrusion, with cancerous cells leaving the epithelium from the membrane side farthest to the TJ domain. Interestingly, our results show SJ-depleted *Ras^V12^* tumors in *Drosophila* leave the flat epithelium apically, also through the cell pole farthest away from SJ. Further studying apical collective delamination in *Drosophila* could help identify conserved molecular regulators involved in vertebrate basal extrusion. Moreover, live delamination is the first step in cancer metastasis and epithelia to mesenchymal (EMT) transition (Yang et al., 2020). Classical EMT hallmarks such as the switch from E-cad to N-cadherin have been revisited. Novel studies show this step is not required for EMT, but rather comprises one of the many possible EMT states (Font-Noguera et al., 2021; Yang et al., 2020). Similarly, initially thought to be a hallmark for EMT, E-cad loss is not required for EMT onset. In some EMT scenarios, E-cad is not lost at all, but instead E-cad remodeling appears tightly regulated across time (Acloque et al., 2024; Campbell & Casanova, 2015; Font-Noguera et al., 2021). Similarly, transcription factors involved in EMT during gastrulation such as Snail and Twist are not implicated in every apical constriction event of EMT, namely single cell delamination during neuroblast ingression (An et al., 2017; Sidor & Röper, 2017; Simões et al., 2017). These observations coincide with our results of persistence of cell-cell adhesion during the collective delamination of living cells via E-cad and β-integrin mediated adhesion. Hence, the cellular steps of collective delamination of living cells identified in this study may occur during EMT onset. Finally, tricellular and bicellular SJ core-components found to synergize with Ras and trigger EMT onset in this study appear frequently mutated in several types of carcinoma. This further cements the role of cell junctions in tumor dissemination (Kyuno et al., 2021).

The experimental findings reported here have direct implications in understanding the cellular steps of the collective delamination of living cells *in vivo*. Moreover, the mathematical framework described here provide insights into how tissue architecture and preferential cell adhesion govern the directionality of delamination, which ultimately determines cell fate, a deeply fundamental question in developmental and cancer biology. Ultimately, we expect our work to help understand how apical and basal collective delamination of living cells differ from each other in terms of junctional remodeling, cell polarity and cell identity, and the underlying mechanism governing directionality in both normal and pathological conditions.

## Materials and methods

### Experimental model

All Drosophila stocks were maintained on media consisting of corn meal, sugar, yeast, and agar on incubators at a constant temperature of 25°C.

### Drosophila genotypes

#### Figure 1

**B, C- *M6^(W18Stop)^*, FRT79E eyMARCM** obtained by crossing yw, eyFlp ; act-Gal4, UAS-nls::GFP ; tub>Gal80, FRT79E / with ; If / CyO ; *M6^(W18Sto)^*, FRT79E / TM6, Tb, Hu

**E, F- UAS-*Ras^V12^*; *M6^(W18Stop)^*, FRT79E eyMARCM** obtained by crossing yw, eyFlp ; act-Gal4, UAS-nls::GFP ; tub>Gal80, FRT79E / with ; UAS-*Ras^V12^* ;*M6^G152)^,* FRT79E / TM6, Tb, Hu

#### Figure 2

**B, C, E, F, H, I- UAS-*Ras^V12^* ; *M6^(W18Stop)^*, FRT79E eyMARCM** obtained by crossing yw, eyFlp ; act-Gal4, UAS-nls::GFP ; tub>Gal80, FRT79E / with ; UAS-*Ras^V12^* ;*M6^G152)^,* FRT79E / TM6, Tb, Hu

#### Figure 3

**B, C, E, F- *aka^L200^*, FRT40A; UAS-*Ras^V12^*; eyMARCM** obtained by crossing *hsFlp,* act-Gal4,UAS-nls::GFP/FM6 ; ptub-Gal80, FRT40A / CyO with ; *aka^L200^*, FRT40A; UAS-*Ras^V12^*/ SM5 – TM6, Tb, Hu

#### Figure 4

**B, D, F, H- UAS-*Ras^V12^* ; *M6^(W18Stop)^*, FRT79E eyMARCM** obtained by crossing yw, eyFlp ; act-Gal4, UAS-GFP ; tub>Gal80, FRT79E / with ; UAS-*Ras^V12^* ;*M6^G152)^,* FRT79E / TM6, Tb, Hu

**J- *aka^L200^*, FRT40A; UAS-*Ras^V12^* ; eyMARCM** obtained by crossing *hsFlp,* act-Gal4,UAS-nls::GFP/FM6 ; ptub-Gal80, FRT40A / CyO with ; *aka^L200^*, FRT40A; UAS-*Ras^V12^* / SM5 – TM6, Tb, Hu

#### Figure 5

**B, E, F, H- UAS-*Ras^V12^* ; *M6^(W18Stop)^*, FRT79E eyMARCM** obtained by crossing yw, eyFlp ; act-Gal4, UAS-GFP ; tub>Gal80, FRT79E / with ; UAS-*Ras^V12^* ;*M6^G152)^,* FRT79E / TM6, Tb, Hu

**C, I- *glil^DV3^*, FRT40A; UAS-*Ras^V12^* ; eyMARCM** obtained by crossing *hsFllp,* act-Gal4,UAS-nls::GFP/FM6 ; ptub-Gal80, FRT40A / CyO with ; *gli^DV^*^3^, FRT40A; UAS-*Ras^V12^* / SM5 – TM6, Tb, Hu

#### Figure 8

**B, C, E, - UAS-*Ras^V12^* ; *M6^(W18Stop)^*, FRT79E eyMARCM** obtained by crossing yw, eyFlp ; act-Gal4, UAS-GFP ; tub>Gal80, FRT79E / with ; UAS-*Ras^V12^* ;*M6^G152)^,* FRT79E / TM6, Tb, Hu

#### Supplementary Figure 1

**B- *M6^(W18Stop)^*, FRT79E eyMARCM** obtained by crossing yw, eyFlp ; act-Gal4, UAS-GFP ; tub>Gal80, FRT79E / with ; If / CyO ; *M6^W18Stop^*, FRT79E / TM6, Tb, Hu

**C, D- *aka^L200^*, FRT40A eyMARCM** obtained by crossing *hsFlp, act*-Gal4, UAS-nls::GFP/FM6 ; ptub-Gal80, FRT40A / CyO with ; *aka^L200^*, FRT40A; MKRS / SM5 – TM6, Tb, Hu

**E, F- *nrv2^K13315^*, FRT40A eyMARCM** obtained by crossing *hsFlp,* act-Gal4,UAS-nls::GFP/FM6 ; ptub-Gal80, FRT40A / CyO with; *nrv2^K13315^*, FRT40A; If / SM5 – TM6, Tb, Hu

#### Supplementary Figure 2

**B,C- UAS-*Ras^V12^*; *M6^(W18Stop)^*, FRT79E eyMARCM** obtained by crossing yw, eyFlp ; act-Gal4, UAS-GFP ; tub>Gal80, FRT79E / with ; UAS-*Ras^V12^* ;*M6^G152)^,* FRT79E / TM6, Tb, Hu

**E, F- *aka^L200^*, FRT40A; UAS-*Ras^V12^* ; eyMARCM** obtained by crossing *hsFlp, act*-Gal4,UAS-nls::GFP/FM6 ; ptub-Gal80, FRT40A / CyO with ; *aka^L200^*, FRT40A; UAS-*Ras^V12^* / SM5 – TM6, Tb, Hu

**G,H- *nrv2^K13315^*, FRT40A; UAS-*Ras^V12^* eyMARCM** obtained by crossing *hsFlp, act*- Gal4,UAS-nls::GFP/FM6 ; ptub-Gal80, FRT40A / CyO with; *nrv2^K13315^*, FRT40A; If / SM5 – TM6, Tb, Hu

#### Supplementary Figure 3

**B, D, F, H, J,- UAS-*Ras^V12^* ; *M6^(W18Stop)^*, FRT79E eyMARCM** obtained by crossing yw, eyFlp ; act-Gal4, UAS-GFP ; tub>Gal80, FRT79E / with ; UAS-*Ras^V12^* ;*M6^G152)^,* FRT79E / TM6, Tb, Hu

**K- *aka^L200^*, FRT40A; UAS-*Ras^V12^* ; eyMARCM** obtained by crossing *hsFlp, act*-Gal4,UAS-nls::GFP/FM6 ; ptub-Gal80, FRT40A / CyO with ; *aka^L200^*, FRT40A; UAS-*Ras^V12^* / SM5 – TM6, Tb, Hu

### Larvae staging

Flies were crossed and flipped ∼48h after initial crossing to stage larvae using as reference the number of hours after egg laying (h AEL). A 2h interval was left after the flip to allow for egg deposition, taking this time point as the “0h AEL” reference.

### Immunostaining and image acquisition

Eye-antenna imaginal discs from staged larvae were dissected and fixed for 15min in 4% formaldehyde & permeabilized/washed in PBT 0.1%, stained with primary antibody solution for 3h at RT, washed and then stained overnight with secondary antibody solution & DAPI (0.5mg/mL, 1:1000 dilution) at 4°C. The antibodies used are detailed in the Key Ressoruces Table.Dissected imaginal discs were mounted in 0,5% N-propylgallate dissolved in 90% glycerol/PBS 1X final & imaged on a Leica inverted confocal SP8 or a Leica upright confocal SPE using a 10X (N.A. 0.3) air objective and oil immersion objectives 20X (N.A. 0.7) & 63X (N.A. 1.4). Images were acquired using a 405 laser to excite DAPI stain and detected with a hybrid detector (HyD). A 488 nm argon laser used to excite the GFP fluorophore, endogenously expressed under the UAS and detected with a HyD. A 561nm DPSS diode laser used to excite Cy-3, detected with a HyD. A 633nm DPSS diode laser used to excite Cy-5 and detected with a photomultiplier tube (PMT4).

### Image processing and analysis

All image processing and analysis was performed using Fiji. For immunofluorescence and fluorescent reporter microscopy-based assays, all measurements were derived from the eye part of the eye-antenna imaginal disc.

### Measuring distance to the morphogenetic furrow

High and low magnification confocal microscopy images were used to measure the distance from the cell in the clone closer to the forefront of the MF, identified at the most posterior apically constricted apex in the tissue indentation of the pseudostratified epithelium. Measurements were carried out for clones in each of the events of tissue fold anterior anterior to the MF (no difference with wild-type counterparts, apical constriction, tissue invagination) and posterior to the MF (apical constriction, tissue invagination) as well as step of apically delaminating clones (octopus, transition in the lumen, clone detachment).

### Quantification of clone size during collective delamination and tissue formation

Confocal microscopy images of *Drosophila* heterozygous eye-antennal imaginal discs immuno-stained with anti-E-cad (grey) and anti-phosphorylated histone 3 (PH3) or Cleaved-Casape-3 (C-Cas3) were analyzed. Clone size was quantified by counting the number of nuclei in the clones of SJ-deficient *RasV^12^* transformed cells, expressing the GFP clone marker. Clone size for each step of apical and basal collective delamination and tissue formation was quantified and average size of clone cluster was obtained.

### Computational modelling

The model has been modified from a previously published model (Plunder et al 2024). Briefly, cells are abstracted to nuclei attached to a set of adjustable springs connecting them to an apical point and a basal point. Apical points of adjacent cells are also attached by springs. Basal points are attached to a simple non-deformable basal line representing the basement membrane. Nuclei repel each other via a repulsion force acting on their soft outer cores. The hard inner cores of nuclei cannot overlap. Loss of apical or basal adhesion (events ***A*** and ***B***) remove the adjustable apical-nuclei or basal-nuclei springs. The new event ***ApC*** increases the strength of the strings between adjacent apical points for clone cells. Parameters were adapted to allow flat and short pseudostratified configurations to emerge. The transition between pseudostratified and flat epithelial epithelium is achieved by changing a limited set of parameters (Supplementary Table 1). For instance, the allowed maximal distance between basal points. When set to 10 μm this distance allows cells to arrange themselves as a flat monolayer. When set to 3 μm it forces cells to redistribute their nuclei along the apicobasal axis to minimize tension/compression and this generates pseudostratification. To prevent the pseudostratified tissue from growing over 3-4 pseudolayers a maximum height is set and acts as a constraint. Since by default the basal compartment located below the basal line is permissive and does not act as a physical obstacle, the flat tissue tends to flip and rotate around the basal line. To prevent this behavior, we added a very weak repulsive within the basal compartment which pushes cells weakly upwards as soon as a nucleus is below the basal line, the strength of this force is 2.5% of the nuclei-nuclei repulsion force strength and does not prevent basal delamination. For simulations with non-permissive matrix we tuned this force to 250 % of the nuclei-nuclei repulsion force. Further modification of the model includes a new event called ***Ai***. During this event, the apical-apical springs at the interface between control cells and clone cells are removed and a recovery phase is added where the two disconnected apical layers of control cells directly connect again with a long apical-apical spring crossing the gap caused by the intermediate EMT cells. The rest length of this apical-apical spring goes to zero with a constant speed of 5μm/h. Montage of simulation outputs into supplementary movies was made using VSDC video editor.

### Key resources table

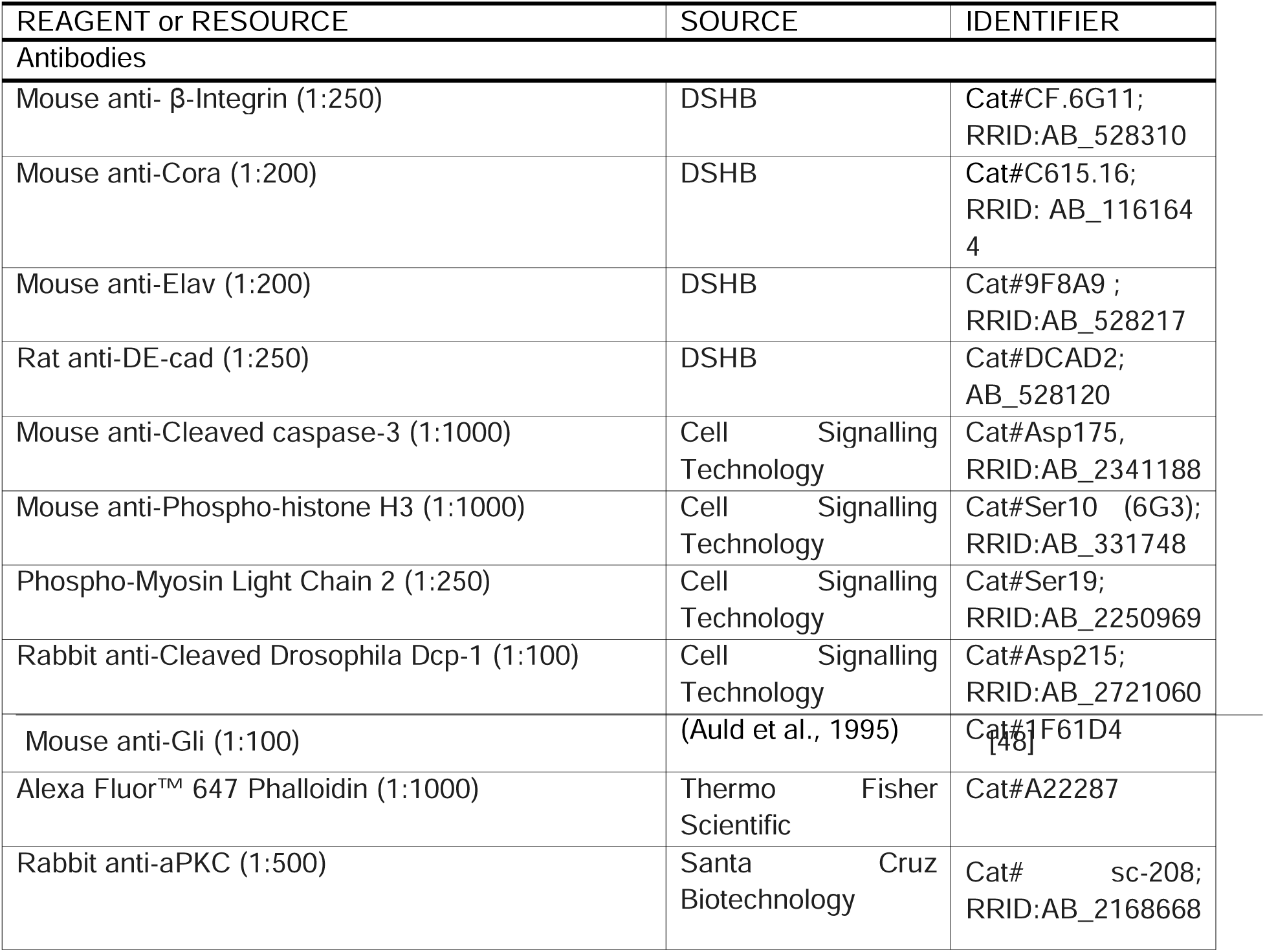
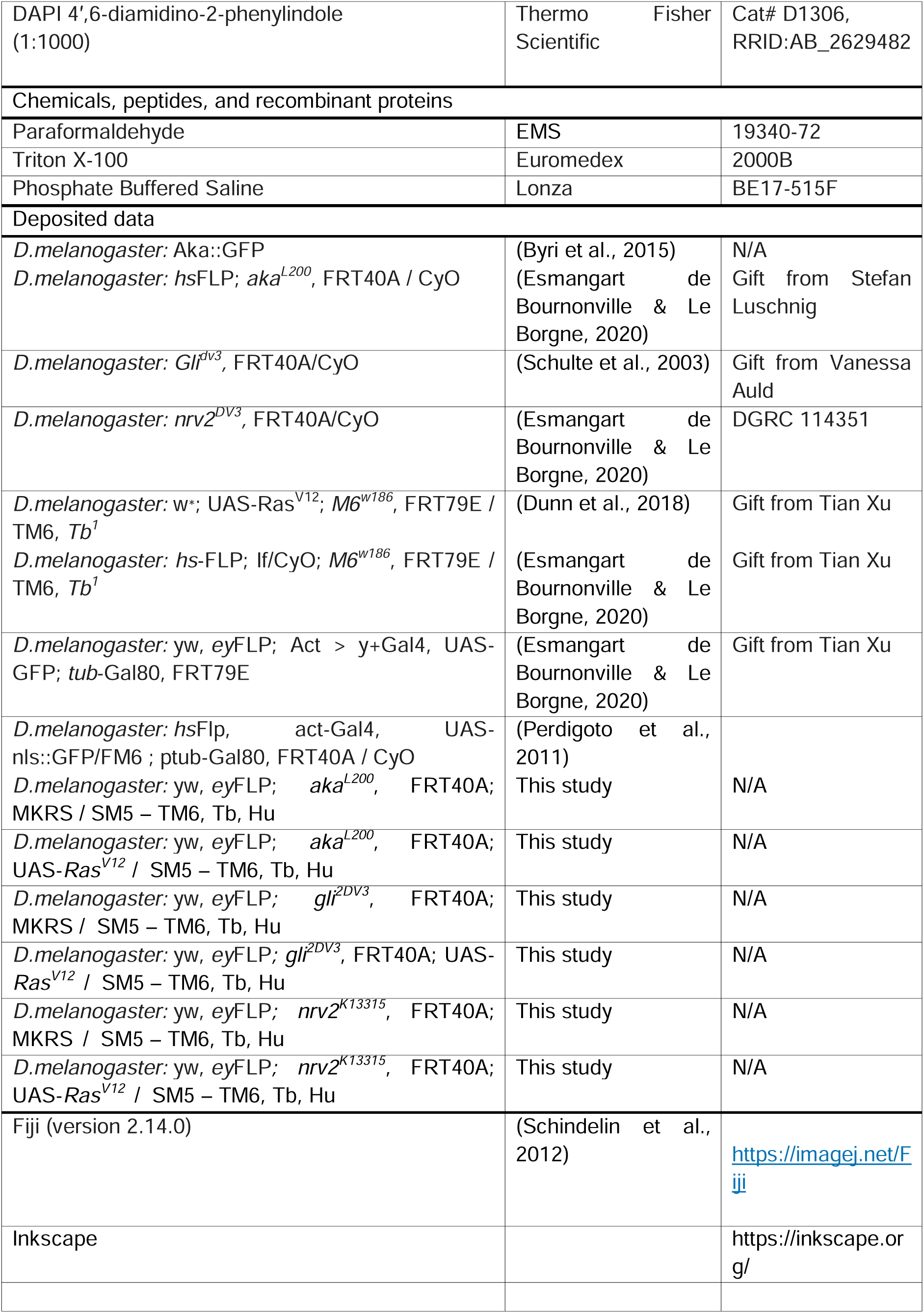
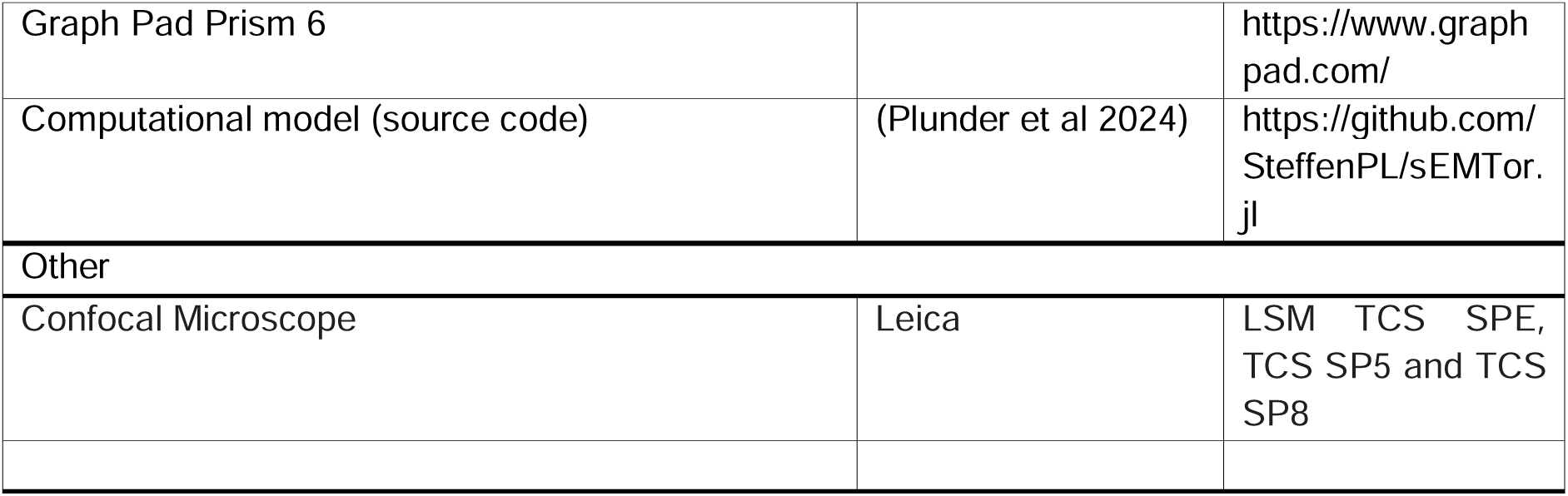

## Acknowledgements

We thank V. Auld, A. Bardin, S. Luschnig, Tian Xu and the Bloomington Stock Center and the National Institute of Genetics Fly Stock Center for providing fly lines. The monoclonal antibodies against Cora, E-cad and β-Integrin were obtained from the Developmental Studies Hybridoma Bank, generated under the auspices of the National Institute of Child Health and Human Development, and maintained by the University of Iowa Department of Biological Sciences. We also thank S Dutertre and X Pinson from the Microscopy Rennes Imaging Center-BIOSIT (France). We are grateful to Chantal Roubinet, Caroline Dillard for critical reading of the manuscript.

---

**Supplementary Figure 1.**
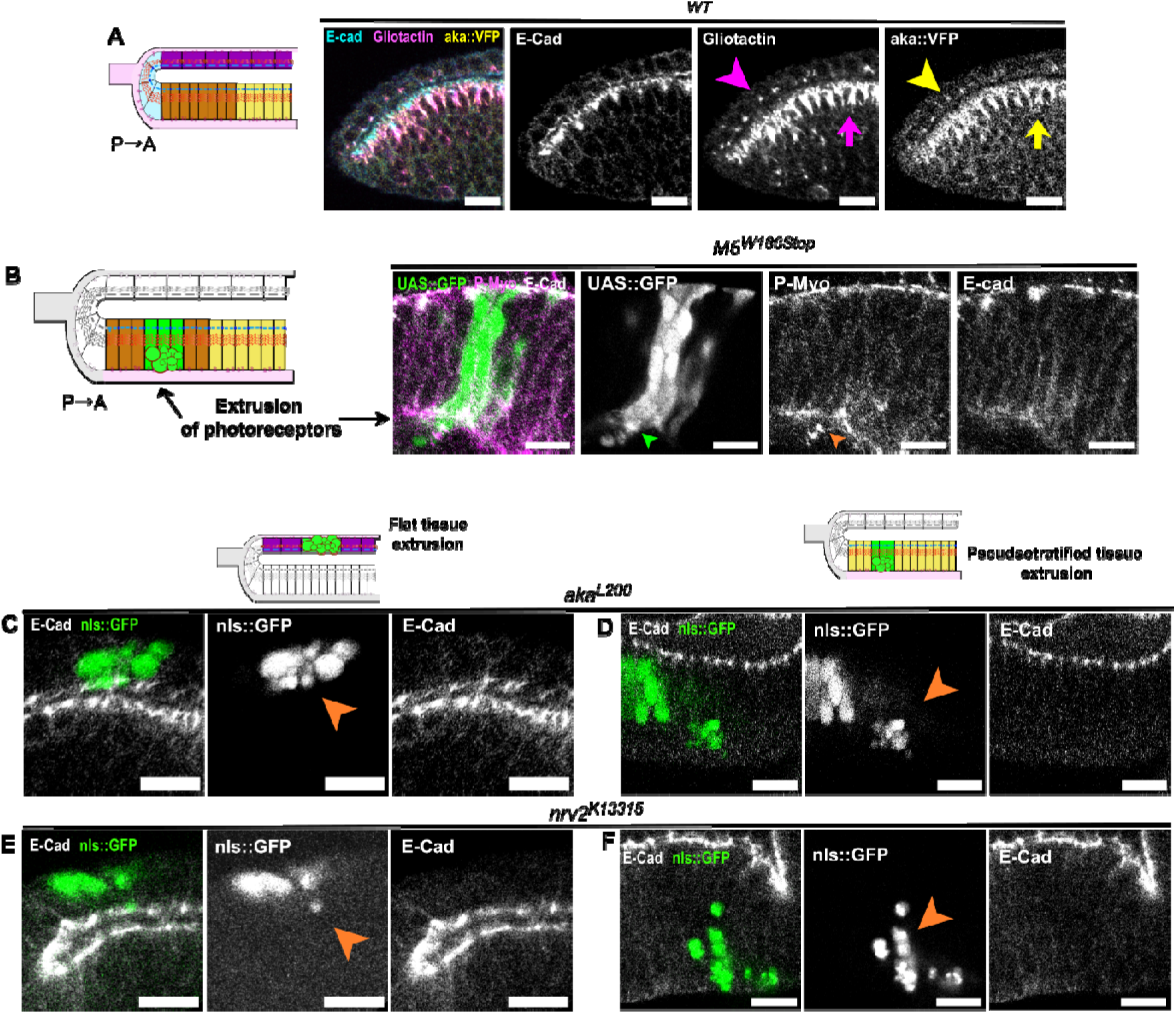
Microscopy images, related to Figures 1 Septate junction loss triggers extrusion in the eye disc. (A, B) Schematic of an orthogonal section along the dorso-ventral axis of the *Drosophila* eye and confocal microscopy images of eye-antennal larval imaginal discs. (A) wild-type eye disc immunostained for E-cad (grey) and Gli (magenta) where all cells express the endogenously tagged aka::VFP (yellow). Cells in the flat epithelium express Gli and aka (pink and yellow arrowheads, respectively), as well as cells in the pseudostratified epithelium (pink and yellow arrows, respectively). (B) Heterozygous eye disc comprised of wild-type and mutant clones tagged with GFP and defective for the tricellular SJ master regulator M6 (*M6^W186Stop^*, green). Mutant cells (green arrowhead) in the differentiated part of the eye disc, posterior to the MF, where photoreceptors and interommatidial cells are found, contain fragmented cytoplasm (green arrowhead) strongly stained with P-Myo (orange arrowhead), a hallmark of apoptotic bodies and cell extrusion. (C, D) Heterozygous eye disc comprised of wild-type and mutant clones tagged with GFP and defective for the tricellular SJ master regulator Aka (*aka^L200^*, green). Mutant cells in the peripodial epithelium (C) and in the pseudostratified epithelium (D) contain pyknotic bodies, suggesting they are eliminated from the tissue via caspase-activated cell extrusion (orange arrowheads). (E, F) Heterozygous eye disc comprised of wild-type and mutant clones tagged with GFP and defective for the bicellular SJ core-complex component Nrv2 (*Nrv2^K3315^*, green). Mutant cells in the peripodial epithelium (E) and in the pseudostratified epithelium (F) pyknotic bodies, suggesting they are eliminated from the tissue via caspase-activated cell extrusion (orange arrowheads). The scale bars represent 10 μm. All images are single confocal optical sections with posterior to the left, anterior to the right. See also Figure 1.

**Supplementary Figure 2.**
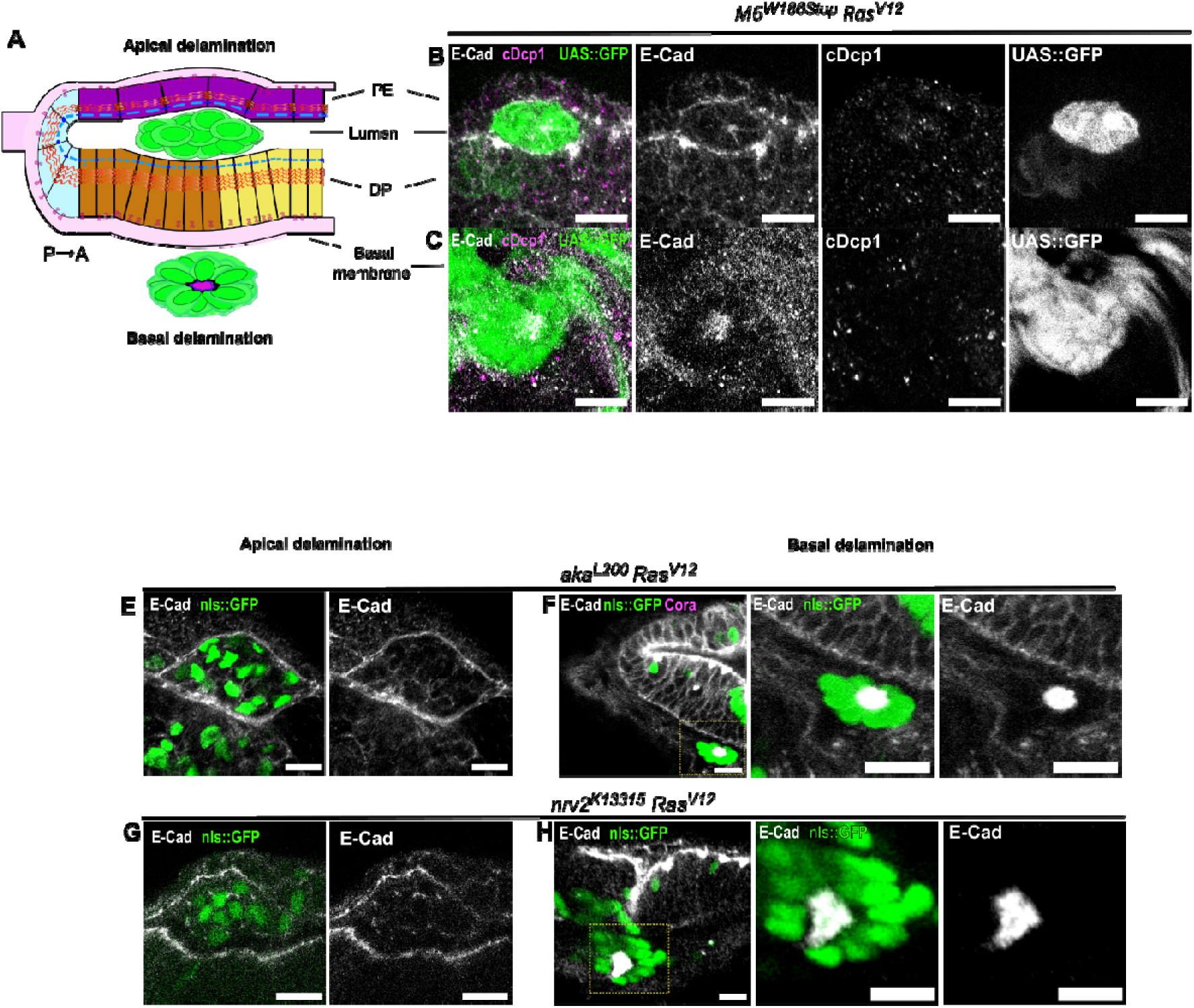
Microscopy images, related to Figures 2 Septate junction loss in synergy with Ras transformation triggers collective delamination in the eye disc. (A) Schematic of an orthogonal section along the dorso-ventral axis of the *Drosophila* eye antennal larval imaginal discs where SJ-deficient RasV12 mutant cells undergo apical and basal collective delamination. (B, C) Heterozygous eye discs immunostained with anti-E-cad (grey) and anti-Drosophila caspase-1 (DCP-1) (C, magenta). Clones tagged with GFP are mutant for M6 and express overactivated Ras (*M6^W186Stop^Ras^V12^*, green). Mutant cells delaminate collectively and alive out of an epithelium, either apically, appearing in the lumen (B), or basally, forming rosettes basal to the eye disc (C). (E,F) Heterozygous eye discs immunostained with anti-E-cad (grey). Clones tagged with GFP are mutant for aka and express overactivated Ras (*aka^L200p^Ras^V12^*, green). Mutant cells delaminate collectively and alive out of an epithelium, either apically, appearing in the lumen (E), or basally, forming rosettes basal to the eye disc (F). (G, H) Heterozygous eye discs immunostained with anti-E-cad (grey). Clones tagged with GFP are mutant for Nrv2 and express overactivated Ras (*nrv2^K13315^Ras^V12^*, green). Mutant cells delaminate collectively and alive out of an epithelium, either apically, appearing in the lumen (G), or basally, forming rosettes basal to the eye disc (H). The scale bars represent 10 μm. All images are single confocal optical sections with posterior to the left, anterior to the right. See also Figure 2.

**Supplementary Figure 3.**
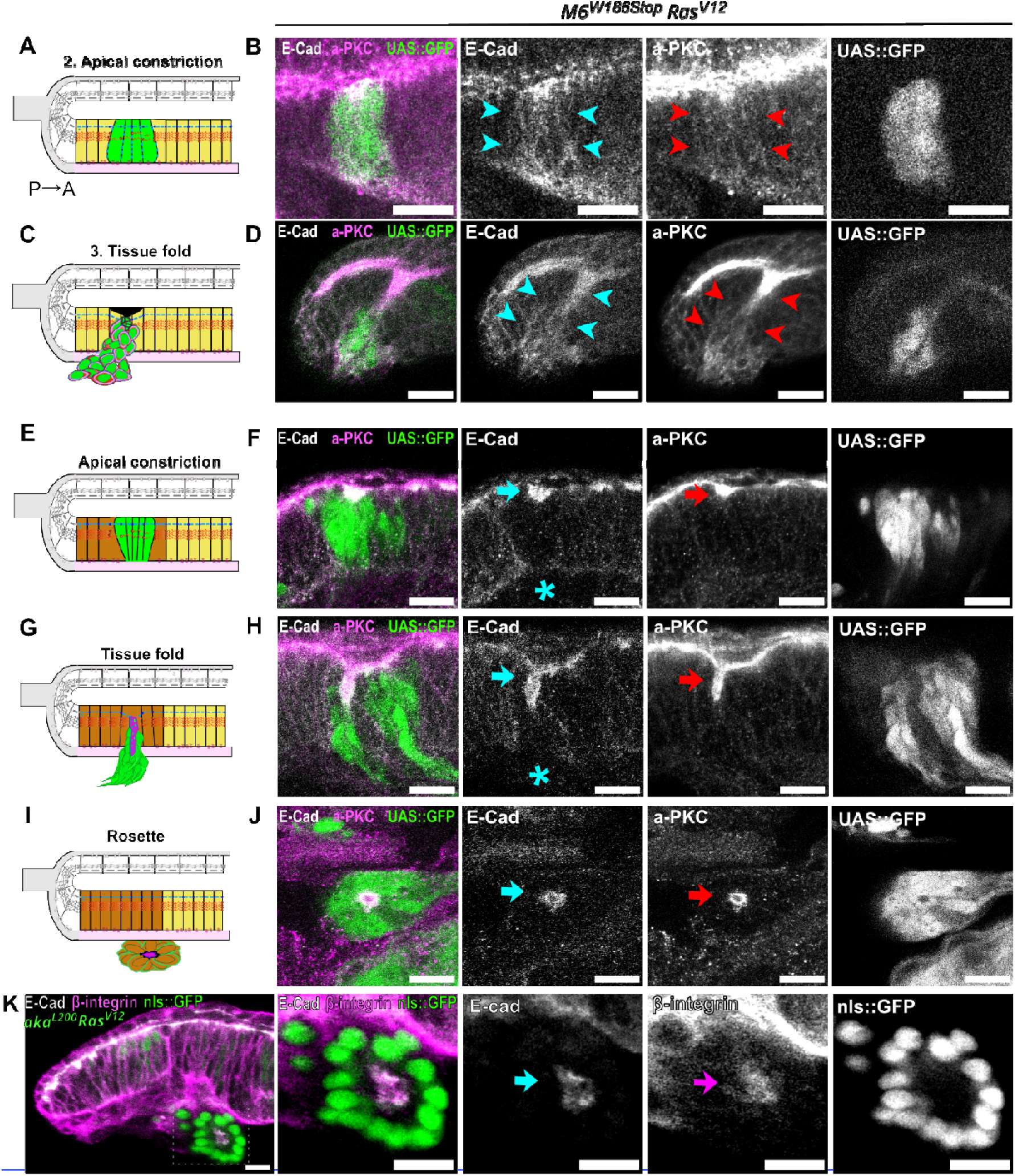
Microscopy images, related to Figures 4. Rosettes are formed independently of tissue folds. (A, C, E, G, I) Schematic of an orthogonal section along the dorso-ventral axis of the *Drosophila* eye antennal larval imaginal discs where SJ-deficient RasV12 mutant cells undergo tissue folding anterior (A, C, yellow) or posterior (E, G, brown) to the MF. Rosettes are formed by differentiated cells (I, brown). (B, D, F, H, J) Heterozygous eye discs immunostained with anti-E-cad (grey) and anti-aPKC (aPKC, magenta). Clones tagged with GFP are mutant for M6 and express overactivated Ras (*M6^W186Stop^Ras^V12^*, green). (B, D) Mutant cells anterior to the MF apically constrict (B) and form tissue folds (D). During both events, basolateral E-cad delocalization (blue arrowheads) is concomitant with a partial loss of polarity, noted by the partial basal localization of aPKC not seen in the wild-type cells (red arrowheads). (F, H) Mutant cells posterior to the MF apically constrict (F) and form tissue folds (H). During both events, E-cad appear well localized apically (blue arrow) and the basal pool of E-cad is lost (blue asterisk). During both events cell polarity is preserved, as noted by the apical restriction of aPKC (red arrows). (J) Rosettes are formed by a polarized cluster of cells organized radially that exhibit Ecad at the center (blue arrow), corresponds to the apical domain, as noted by aPKC restricted (red arrow). (K) Heterozygous eye discs immunostained with anti-E-cad (grey) and anti-β-integrin (magenta). Clones tagged with GFP are mutant for aka and express overactivated Ras (aka*^L200^Ras^V12^*, green) Rosettes exhibit adhesion proteins at the center of the radial cluster, zt the apical domain, namely E-cad (blue arrow) and β-integrin (pink arrow). The scale bars represent 10 μm. All images are single confocal optical sections with posterior to the left, anterior to the right. See also Figure 4.

**Supplementary Figure S4.**
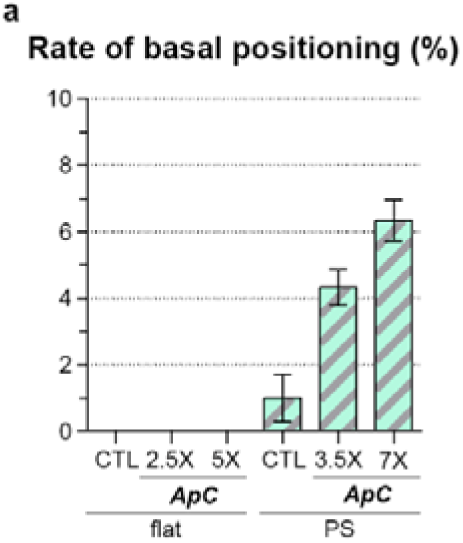
Apical constriction does not lead to delamination but is sufficient to trigger basal displacement of cells in the pseudostratified epithelium. (A) Mean rate of basal positioning with standard error of the mean. To assess the effect of apical constriction, apical contractility of clone cells was increased 2.5 times (2.5X) and 5 times (5X) the control level in the flat epithelium and 3.5 times (3.5X) and 7 times (7X) the control level in the PS epithelium. ApC, apical constriction; CTL, control levels of apical contractility; PS, pseudostratified tissue.

**Figure S5.**
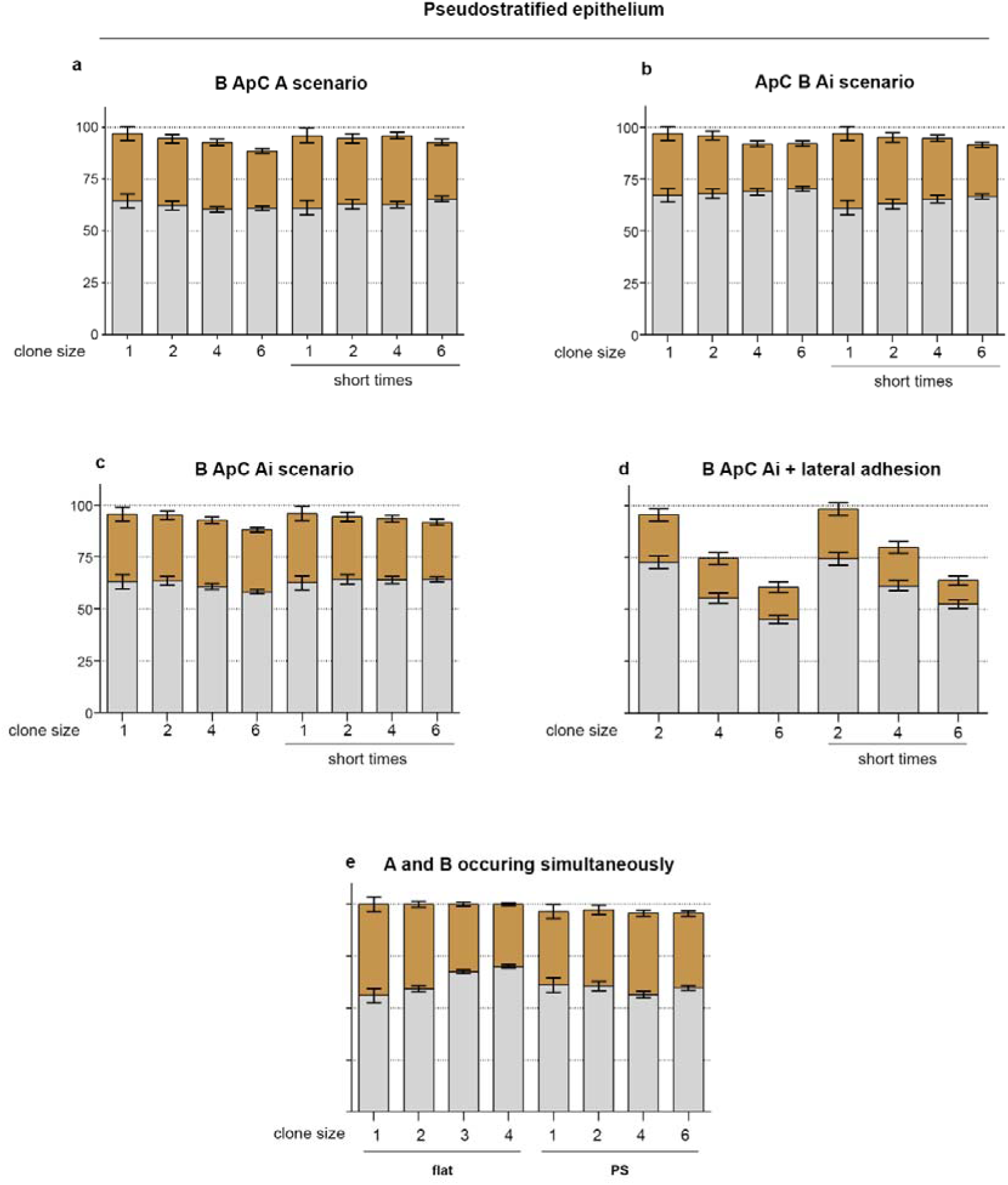
Timing of epithelial destabilization has marginal influence on the directionality of delamination. (A-D) Mean rates of delamination with standard error of the mean for all indicated scenarios tested in clones of various sizes in the peusdostratified epithelium. Scenarios have a time window of 20 hours to be implemented whereas under the “short times” condition all events must occur within 4 hours. (E) Mean rates of delamination with standard error of the mean when events and A and B occur simultaneously. Each simulation was performed 200 times. A, loss of apical adhesion between clone cells and between clone cells and control cells; Ai, loss of apical adhesion only at the interface between clone cells and control cells; ApC, apical constriction; B, loss of basal adhesion; LA, lateral adhesion; PS, pseudostratified.

## Supplementary Movies legends

**Supplementary Movie 1. Computational modelling of the eye-antennal imaginal disc using the EMT-simulator**

Initialization of each type of epithelium. All cells are control cells. Top panel: flat tissue, bottom panel: short pseudostratified (PS) tissue. Apical points are in red, basal points in black. Large nuclei correspond to mitosis. The flat epithelium reaches its stable configuration in about 5 hours. The pseudostratification with 2 to 3 pseudolayers takes about 20 hours to emerge and the stable configuration is reached at around 30 hours. In the PS tissue cells undergo interkinetic movement. For instance, the central cell labelled in magenta divides at 9, 23 and 34 during the simulation. Total duration 40 hours, time stamp in the top right corner.

**Supplementary Movie 2. The outcome of epithelial destabilization is sensitive to clone size in flat but not in pseudostratified epithelia**

Simulation of the ApC A B scenario in flat (top panel) and pseudostratified tissues (bottom panels) in a single cell (left panels) or a clone (right panels). ApC is implemented since the onset of simulation, A occurs at 40h, B occurs at 50h. Movie starts at 30h post initiation and stops at 80h of simulation.

**Supplementary Movie 3. Moderate cell adhesion does not prevent clone splitting in the pseudostratified epithelium but reverses directionality in the flat layer**

Simulation of the ApC Ai B scenario in flat (top panel) and pseudostratified tissues (bottom panels. ApC is implemented since the onset of simulation, Ai occurs at 40h, B occurs at 50h. Movie starts at 30h post initiation and stops at 80h of simulation.

**Supplementary Movie 4. Reinforced lateral adhesion between clones in pseudostratified epithelia suppresses clone splitting and favours basal delamination** Simulation of the ApC Ai B and lateral adhesion between clone cells scenario in flat (top panel) and pseudostratified tissues (bottom panels). ApC is implemented since the onset of simulation, Ai occurs at 40h, B occurs at 50h. Movie starts at 30h post initiation and stops at 80h of simulation.

**Supplementary Movie 5. A non-permissive matrix prevents basal delamination in flat epithelia but does not abrogate the intrinsic basal directionality of delamination upon epithelial destabilization in the pseudostratified layer**

Simulation of the ApC Ai B and lateral adhesion between clone cells scenario in flat and pseudostratified tissues (top panels). Simulation of the ApC B and lateral adhesion between clone cells scenario in flat and pseudostratified tissues (middle panels). Simulation of the B ApC and lateral adhesion between clone cells scenario in flat and pseudostratified tissues (bottom panels). ApC is implemented since the onset of simulation for the ApC Ai B and ApC B scenario and implemented at 50 hours for the B ApC scenario. Ai occurs at 40h, B occurs at 50h. Movie starts at 30h post initiation and stops at 80h of simulation.

